# Kinetic improvement of an algal diacylglycerol acyltransferase 1 via fusion with an acyl-CoA binding protein

**DOI:** 10.1101/2019.12.21.885871

**Authors:** Yang Xu, Kristian Mark P. Caldo, Lucas Falarz, Kethmi Jayawardhane, Guanqun Chen

**Affiliations:** Department of Agricultural, Food and Nutritional Science, University of Alberta, Edmonton, Alberta, Canada T6G 2P5; Department of Biological Sciences, University of Manitoba, Winnipeg, Manitoba, Canada R3T 2N2

**Keywords:** Triacylglycerol biosynthesis, DGAT, acyl-CoA binding protein, algal lipid, enzyme kinetics, tobacco, yeast, *Chromochloris zofingiensis*

## Abstract

Microalgal oils in the form of triacylglycerols (TAGs) are broadly used as nutritional supplements and biofuels. Diacylglycerol acyltransferase (DGAT) catalyzes the final step of acyl-CoA-dependent biosynthesis of TAG and is considered a key target for manipulating oil production. Although a growing number of *DGAT1*s have been identified and over-expressed in some algal species, the detailed structure-function relationship, as well as the improvement of DGAT1 performance via protein engineering, remain largely untapped. Here, we explored the structure-function features of the hydrophilic N-terminal domain of DGAT1 from the green microalga *Chromochloris zofingiensis* (CzDGAT1). The results indicated that the N-terminal domain of CzDGAT1 was less disordered than those of the higher eukaryotic enzymes and its partial truncation or complete removal could substantially decrease enzyme activity, suggesting its possible role in maintaining enzyme performance. Although the N-terminal domains of animal and plant DGAT1s were previously found to bind acyl-CoAs, replacement of CzDGAT1 N-terminus by an acyl-CoA binding protein (ACBP) could not restore enzyme activity. Interestingly, the fusion of ACBP to the N-terminus of the full-length CzDGAT1 could enhance the enzyme affinity for acyl-CoAs and augment protein accumulation levels, which ultimately drove oil accumulation in yeast cells and tobacco leaves to higher levels than the full-length CzDGAT1. Overall, our findings unravel the distinct features of the N-terminus of algal DGAT1 and provide a strategy to engineer enhanced performance in DGAT1 via protein fusion, which may open a vista in generating improved membrane-bound acyl-CoA-dependent enzymes and boosting oil biosynthesis in plants and oleaginous microorganisms.

**SIGNIFICANCE STATEMENT:** Here, we explored the N-terminus of a microalgal DGAT1, a membrane-bound enzyme determining oil biosynthesis, using *in silico* analysis, truncation mutagenesis, protein fusion and *in vitro* and *in vivo* characterization, and demonstrated its distinct structure-function features from the higher eukaryotic enzymes. We further engineered enhanced performance in DGAT1 via N-terminal fusion of ACBP, and obtained a kinetically improved enzyme with augmented protein production levels, which could boost oil accumulation in yeast and plant vegetative tissues.

## INTRODUCTION

Plant-derived triacylglycerol (TAG) is one of the most abundant forms of energy storage and reduced carbon in nature, which has been widely used as food, feed and renewable feedstocks for industrial applications (Xu, Caldo, *et al*., 2018). Microalgae hold the promise of a sustainable bioresource of TAG because of the high ability to accumulate lipids and less competition for arable land with food crops (Xu, Caldo, *et al*., 2018; Hu *et al*., 2008). In recent years, the emerging research interest in exploring the oil biosynthesis mechanisms in microalgae has opened an important vista to fulfil the potential of microalgal oil production via physiological and genetic manipulations.

In microalgae, TAG assembly typically occurs through acyl-CoA-dependent and acyl-CoA-independent pathways by a series of acyltransferases which are universally present in plants and animals (Xu, Caldo, *et al*., 2018; Kong *et al*., 2018). Among them, acyl-CoA:diacylglycerol acyltransferase (DGAT, EC 2.3.1.20) catalyzes the acylation of *sn*-1,2-diacylglycerol with acyl-CoA to produce TAG, which is the final committed step in the acyl-CoA-dependent TAG biosynthesis. DGAT appears to play a prominent role in affecting the flux of carbon into TAG in many oilseed crops (Liu *et al*., 2012; Katavic *et al*., 1995; Zou *et al*., 1999; Weselake *et al*., 2008) and has been regarded as an important target for manipulation. Two major forms of membrane-bound non-homologous DGAT, designated DGAT1 and DGAT2, are known to predominantly contribute to TAG formation in developing seeds and microalgae. In plants, DGAT1 is considered as a major player in seed oil accumulation in some oil crops, such as rapeseed (*Brassica napus*) and safflower (*Carthamus tinctorius*) (Rahman *et al*., 2013; Tzen *et al*., 1993; Weselake *et al*., 1993), whereas DGAT2 appears to play a minor role in affecting oil production in oil crops. DGAT2, however, is important for incorporating unusual fatty acids into storage TAG in plants, such as tung tree (*Vernicia fordii*), castor (*Ricinus communis*), and ironweed (*Vernonia galamensis*) (Shockey *et al*., 2006; Kroon *et al*., 2006; Li *et al*., 2010). In microalgae, on the other hand, one or two copies of *DGAT1* and several copies of *DGAT2* were found to likely contribute to the complexity of TAG formation, although their physiological roles remain ambiguous (Mao *et al*., 2019; Chen and Smith, 2012; Gong *et al*., 2013; Liu and Benning, 2013; Xu, Caldo, *et al*., 2018; Turchetto-Zolet *et al*., 2011; Liu *et al*., 2016). Given the importance of the enzyme in governing the flux of substrates into TAG, over-expression of *DGAT* cDNAs have been used to manipulate oil production in the seeds of *Arabidopsis thaliana* and oilseed crops such as soybean (*Glycine max*), *B. napus*, corn (*Zea mays*) and *Camelina sativa* (Jako *et al*., 2001; Weselake *et al*., 2008; Roesler *et al*., 2016; Kim *et al*., 2016; Oakes *et al*., 2011; Lardizabal *et al*., 2008; Li *et al*., 2012), in the leaves of *Nicotiana tabacum*, *N. benthamiana*, and *Z. mays* (Alameldin *et al*., 2017; Bouvier-Navé *et al*., 2000; Vanhercke *et al*., 2017; Chen *et al*., 2017), and in oleaginous yeast (Greer *et al*., 2015; Chen *et al*., 2017) and several microalgae including *Chlamydomonas reinhardtii*, *Phaeodactylum tricornutum*, and *Nannochloropsis oceanica* (Zulu *et al*., 2017; Xin *et al*., 2017; Xin *et al*., 2018; Mao *et al*., 2019; Iwai *et al*., 2014).

DGAT1 is an integral membrane protein with multiple transmembrane domains (TMD), the three-dimensional structure of which has not yet been elucidated (Xu, Caldo, *et al*., 2018). DGAT1 shares common features among different organisms, containing a very variable hydrophilic N-terminus with possibly distinct functions and a conserved C-terminal region with 8-10 predicted TMD (Liu *et al*., 2012). The hydrophilic N-termini of *B. napus* and mouse (*Mus musculus*) DGAT1s have been found to be involved in acyl-CoA binding and self-association (Weselake *et al*., 2006; Siloto *et al*., 2008; McFie *et al*., 2010). Recently, the structure of the N-terminal domain of *B. napus* DGAT1 was solved, revealing its important role as an enzyme regulatory domain that positively and negatively modulates enzyme activity (Caldo *et al*., 2017). The N-terminal domain of *B. napus* DGAT1 consists of two different segments, an intrinsically disordered region encompassing an autoinhibitory motif and a folded segment containing the allosteric site for acyl-CoA and CoA for activation and feedback inhibition of the enzyme, respectively (Caldo *et al*., 2017). Although DGAT1s have been characterized from a growing number of microalgal species (Kirchner *et al*., 2016; Wei *et al*., 2017; Guo *et al*., 2017; Guihéneuf *et al*., 2011), the structure-function features of algal DGAT1, as well as using the knowledge in improving DGAT performance, remain largely untapped.

The aim of this study, therefore, is to use a DGAT1 from *Chromochloris zofingiensis* (CzDGAT1), an emerging model green microalgal species for studying TAG and secondary carotenoid accumulation and industrial production, to explore the structure and function features of the N-terminal domain of green microalgal DGAT1 and to investigate the potential of protein fusion in improving DGAT1 performance. After comparing the evolutionary and structural features of algal DGAT1 with the higher eukaryotic enzymes, the function of the hydrophilic N-terminal domain of CzDGAT1 was examined via truncation mutagenesis, protein fusion and *in vitro* enzyme assay. This N-terminus was found to have very different features from those of the plant and animal enzymes and is important for maintaining high DGAT1 activity but not essential for catalysis. The subsequent fusion of an *A. thaliana* acyl-CoA binding protein (AtACBP6) to the N-terminus of CzDGAT1 resulted in a kinetically improved enzyme with augmented protein production levels, and this improved DGAT1 variant could drive oil accumulation to higher levels than the native DGAT1in yeast cells and *N. benthamiana* leaves. The results indicated the fusion of ACBP with DGAT1 may represents a promising strategy in engineering oil production in oleaginous organisms, which may also be used in engineering other membrane-bound acyl-CoA-dependent enzymes.

## RESULTS

### CzDGAT1 is phylogenetically related to plant DGAT1

Phylogenetic analysis was carried out with DGAT1s from *C. zofingiensis* and other algae, plants and animals. The two reported full-length *CzDGAT1* sequences, sharing 39.9% amino acid pairwise identity, were both used in this analysis (Roth *et al*., 2017; Mao *et al*., 2019). As shown in Figure 1, the results revealed some interesting features from an evolutionary perspective. DGAT1s were found to be separated into four subgroups, with animal and plant DGAT1 falling into two separate groups. DGAT1 from the charophyte green alga *Klebsormidium nitens* is grouped with plant DGAT1, whereas CzDGAT1 and other DGAT1 from chlorophyte green algae form a separate group, which is closely related to the plant DGAT1 group. Diatom DGAT1, on the other hand, is clustered with fungal DGAT1 and is separate from all other sequences.

**Figure 1.**
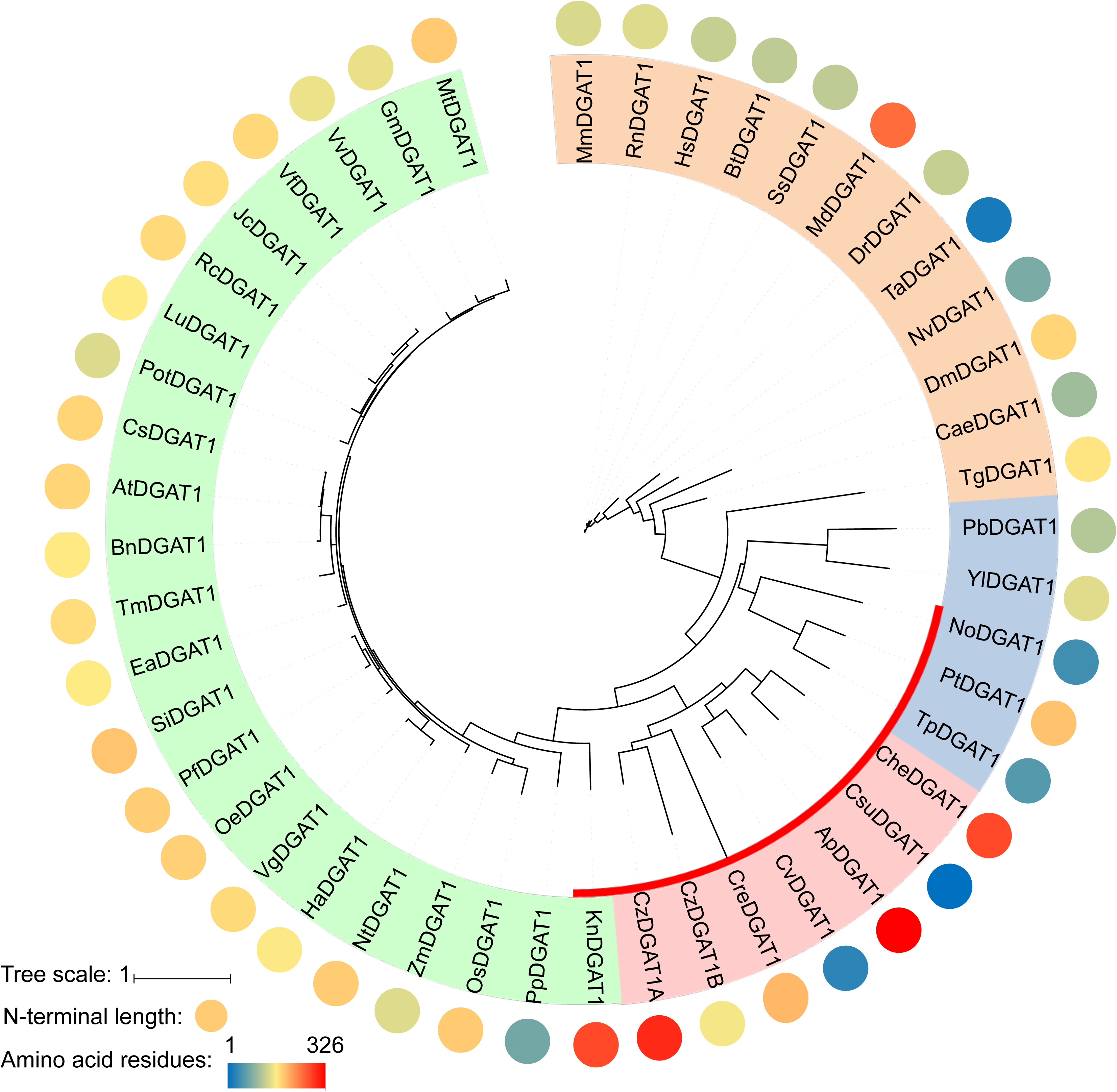
Phylogenetic relationship among CzDGAT1 and DGAT1 from other organisms. The organism and Phytozome/GenBank accession number/JGI protein ID for each protein sequence are shown as follows: *Auxenochlorella protothecoides*, *Ap*, ApDGAT1 (XP_011402032); *Arabidopsis thaliana*, *At*, AtDGAT1 (NM_127503); *Brassica napus*, *Bn*, BnDGAT1 (JN224473); *Bos taurus*, *Bt*, BtDGAT1 (AAL49962); *Caenorhabditis elegans*, *Cae*, CaeDGAT1 (NM_001269372); *Chlorella ellipsoidea*, *Che*, CheDGAT1 (KT779429); *Chlamydomonas reinhardtii*, *Cre*, CreDGAT1 (Cre01.g045903); *Camelina sativa*, *Cs*, CsDGAT1 (XM_010417066); *Coccomyxa subellipsoidea* C-169, *Csu*, CsuDGAT1 (54084); *Chlorella vulgaris*, *Cv*, CvDGAT1 (ALP13863); *Chromochloris zofingiensis*, *Cz*, CzDGAT1A (MH523419), CzDGAT1B (Cz09g08290); *Drosophila melanogaster*, *Dm*, DmDGAT1 (AF468649); *Danio rerio*, *Dr*, DrDGAT1 (NM_199730); *Euonymus alatus*, *Ea*, EaDGAT1 (AY751297); *Glycine max*, *Gm*, GmDGAT1 (AY496439); *Helianthus annuus*, *Ha*, HaDGAT1 (HM015632); *Homo sapiens*, *Hs*, HsDGAT1 (NM_012079); *Jatropha curcas*, *Jc*, JcDGAT1 (DQ278448); *Klebsormidium nitens*, *Kn*, KnDGAT1 (GAQ91878); *Linum usitatissimum*, *Lu*, LuDGAT1 (KC485337); *Monodelphis domestica*, *Md*, MdDGAT1 (XM_007488766); *Mus musculus*, *Mm*, MmDGAT1 (AF078752); *Medicago truncatula*, *Mt*, MtDGAT1 (XM_003595183); *Nannochloropsis oceanica*, *No*, NoDGAT1 (KY073295); *Nicotiana tabacum*, *Nt*, NtDGAT1 (AF129003); *Nematostella vectensis*, *Nv*, NvDGAT1 (XM_001639301); *Olea europae*, *Oe*, OeDGAT1 (AY445635); *Oryza sativa*, *Os*, OsDGAT1 (NM_001061404); *Paracoccidioides brasiliensis*, *Pb*, PbDGAT1 (EEH17170); *Perilla frutescens*, *Pf*, PfDGAT1 (AF298815); *Populus trichocarpa*, *Pot*, PotDGAT1 (XM_006371934); *Physcomitrella patens*, *Pp*, PpDGAT1 (XM_001770877); *Phaeodactylum tricornutum*, *Pt*, PtDGAT1 (HQ589265); *Ricinus communis*, *Rc*, RcDGAT1 (NM_001323734); *Rattus norvegicus*, *Rn*, RnDGAT1 (AB062759); *Sesamum indicum*, *Si*, SiDGAT1 (JF499689); *Sus scrofa*, *Ss*, SsDGAT1 (NM_214051); *Trichoplax adhaerens*, *Ta*, TaDGAT1 (XM_002111989); *Toxoplasma gondii*, *Tg*, TgDGAT1 (AY327327); *Tropaeolum majus*, *Tm*, TmDGAT1 (AY084052); *Thalassiosira pseudonana*, *Tp*, TpDGAT1 (XM_002287179); *Vernicia fordii*, *Vf*, VfDGAT1 (DQ356680); *Vernonia galamensis*, *Vg*, VgDGAT1 (EF653276); *Vitis vinifera*, *Vv*, VvDGAT1 (CAN80418); *Yarrowia lipolytica*, *Yl*, YlDGAT1 (XM_502557); and *Zea mays*, *Zm*, ZmDGAT1 (EU039830). DGAT1s from the animal, fungal, green algal, and plant groups are shown in orange, blue, pink and green, respectively. Algal DGAT1s are shown by red bars. The length of the N-terminus of DGAT1 is shown as a heat map (circle).

Further sequence analysis revealed that similar to plant and animal DGAT1s, the C-terminal portion of algal DGAT1s contains 7-10 predicted transmembrane domains and is the most conserved region, whereas the hydrophilic N-terminus preceding the first predicted transmembrane domain is less conserved and variable in length (Figures 1 and S1 and Table S1). Plant and animal DGAT1s have a hydrophilic N-terminus with a length of approximately 110 and 94 amino acid residues, respectively. On the contrary, the N-terminus of the algal DGAT1 has a quite variable length, ranging from 20 (*Chlorella vulgaris* DGAT1) to 326 amino acid residues (*Auxenochlorella protothecoides* DGAT1). One CzDGAT1 isoform (isoform CzDGAT1B in Mao *et al*., 2019) has a N-terminus composed of 107 amino acid residues (Table S1), which is similar to that of plant and animal DGAT1, and was used for subsequent experiments.

### CzDGAT1 encodes an active enzyme and has a hydrophilic N-terminus with less propensity to become disordered

The functionality of CzDGAT1 was characterized using the yeast mutant H1246, which is devoid of TAG biosynthesis ability (Sandager *et al*., 2002). The yeast complementation and fatty acid feeding assays showed that CzDGAT1 was able to restore TAG biosynthesis in *Saccharomyces cerevisiae* mutant H1246 (Figure 2A) and facilitated the incorporation of the exogenously fed linoleic acid (C18:2*Δ^9cis,12cis^*, C18:2) and *α*-linolenic acid (C18:3*Δ^9cis, 12cis, 15cis^*, C18:3) into yeast TAG (Figure 2B). Further *in vitro* enzyme assay confirmed that CzDGAT1 displayed a strong DGAT activity (Figure 2C).

**Figure 2.**
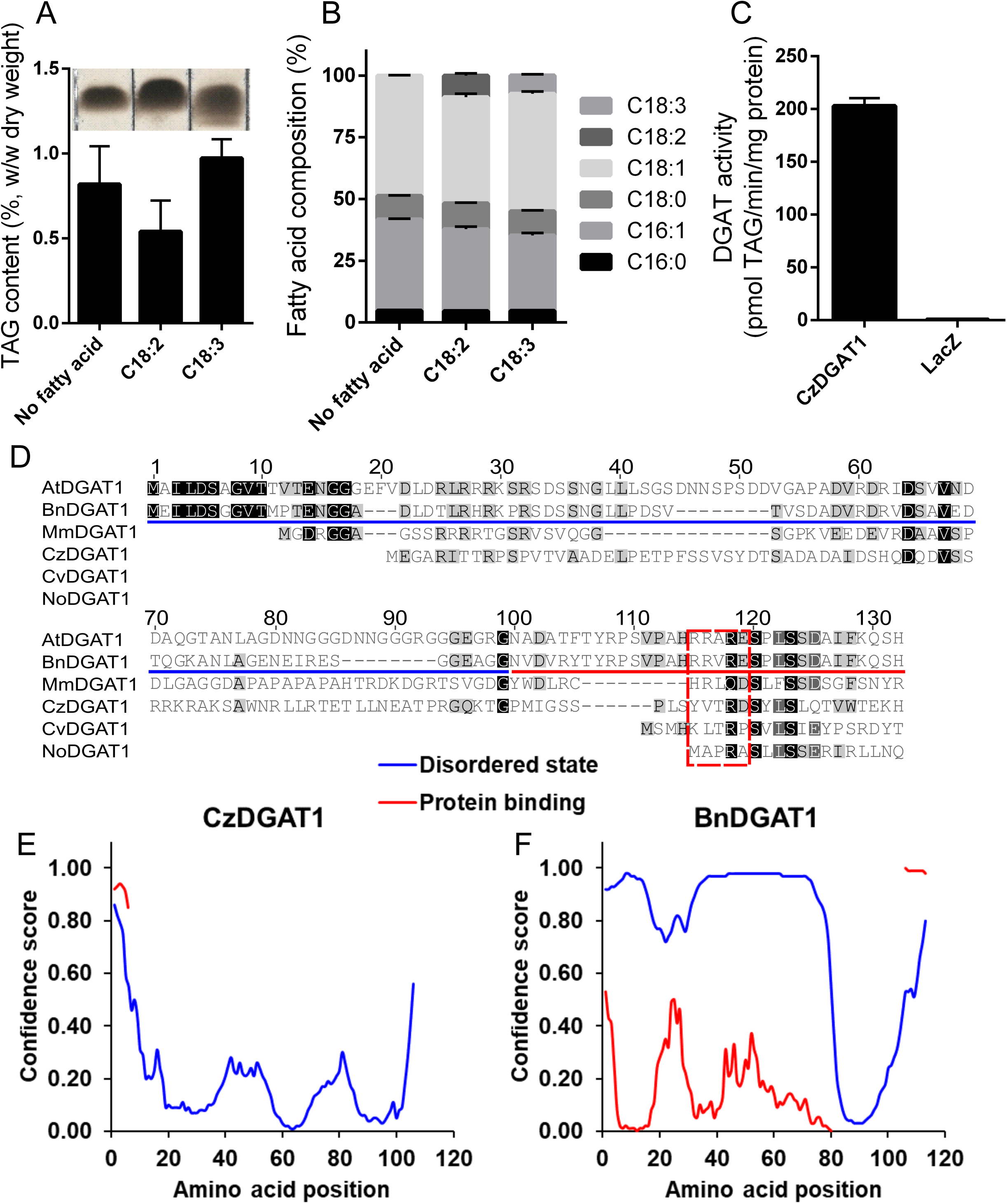
CzDGAT1 encodes an active enzyme and has a unique N-terminus with less propensity to become disordered. A and B, Triacylglycerol (TAG) content (A) and fatty acid (FA) composition (B) in yeast producing CzDGAT1 cultured in the absence of FA or the presence of exogenous linoleic acid (18:2*Δ^9cis,12cis^*, 18:2), or α-linolenic acid (C18:3*Δ^9cis,12cis,15cis^*, 18:3). Yeast cells were harvested after 48 h of induction for lipid analysis. C16:0, Palmitic acid; C18:0, Stearic acid; C18:1, Oleic acid. C, *In vitro* DGAT assay using yeast microsomal fractions containing recombinant CzDGAT1. D, Sequence alignment of the N-terminal regions of DGAT1. *At*, *Arabidopsis thaliana*; *Bn*, *Brassica napus*; *Cv*, *Chlorella vulgaris*; *Cz*, *Chromochloris zofingiensis*; *No*, *Nannochloropsis oceanica*. E and F, Prediction of intrinsic disorder profile (blue) of the N-terminal regions of CzDGAT1 (D) and BnDGAT1 (E) and likelihood to participate in protein-protein interaction (red) by DISOPRED analysis (Ward *et al*., 2004). For A, B and C, data represent means ± S.D. (n = 3).

The N-terminal region of plant DGAT1 has been found to serve important regulatory functions (Caldo *et al*., 2017). The majority of the N-terminal hydrophilic region of plant and animal DGAT1 is likely present in a disordered state, whereas only a small portion preceding the first predicted transmembrane domain appears to have secondary structure (Figure 2D) (Caldo *et al*., 2017). To explore whether algal DGAT1 has similar features, the secondary structure of the N-terminal region of algal DGAT1 was analyzed using DISOPRED (Ward *et al*., 2004). Interestingly, CzDGAT1 may have a very different profile in the N-terminus, where there is much less propensity to be disordered than that of *B. napus* DGAT1 (Figures 2E and F). Similarly, N-terminus with less disordered state was also predicted in DGAT1 from a few other algal species (Figure S2). On the other hand, algal DGAT1 with an extremely long N-terminus, such as DGAT1 from *K. nitens*, is predicted to have both disordered and less disordered N-terminal segments. Furthermore, the folded portion of the N-terminal region of DGAT1 has been found to contain an allosteric site for binding of acyl-CoA and/or CoA has been identified in *B. napus* DGAT1 (Caldo *et al*., 2017). To test whether the N-terminus of algal DGAT1 also has the regulatory features, the N-terminal regions of DGAT1 from several plants, animal and algal species were aligned (Figure 2D). The four amino acid residues (R96, R97, R99 and E100 in *B. napus* DGAT1) implicated in CoA binding (Caldo *et al*., 2017) appear to be highly conserved in plant and animal DGAT1 but not in algal DGAT1, in which only the third residue (R99) is conserved. It should be noted that CzDGAT1 contains partial of the allosteric site in its N-terminus, where two out of the four amino acid residues involved in CoA binding are conserved (Figure 2D).

### N-terminal truncation of CzDGAT1 leads to less active enzymes

Considering the partial conservation of the allosteric sites in the N-terminus of CzDGAT1, it is interesting to test whether the N-terminus of CzDGAT1 also serves an analogous function to the *B. napus* DGAT1 N-terminus in catalysis. CzDGAT1 is predicted to have a 107-amino acid residue-long hydrophilic N-terminal region, followed by 9 predicted hydrophobic segments (Figure 3A). To probe the possible role of the N-terminal region, the full-length CzDGAT1 and two N-terminal truncated versions were produced in *S. cerevisiae* mutant H1246 and the microsomal fractions containing the recombinant proteins were used to determine DGAT activity and protein production levels. Both the removal of the first 80 amino acid residues (CzDGAT1_81_-_550_), which roughly correspond to the intrinsically disordered region in the N-terminus of *B. napus* DGAT1 (Figure 2D), and the entire N-terminal region (CzDGAT1_107-550_) led to reduced enzyme production levels (Figure 3B) and enzyme specific activities (Figure 3C). The specific activity of each enzyme was then normalized by the corresponding protein production level and the normalized activities of CzDGAT1_81-550_ and CzDGAT1_107-550_ were about 10 and 120-fold lower than that of the full-length enzyme, respectively (Figure 3D). These results suggest that the first 80 amino acids are not dispensable for the enzyme activity, and the entire N-terminal domain may be important for maintaining high enzyme activity.

**Figure 3.**
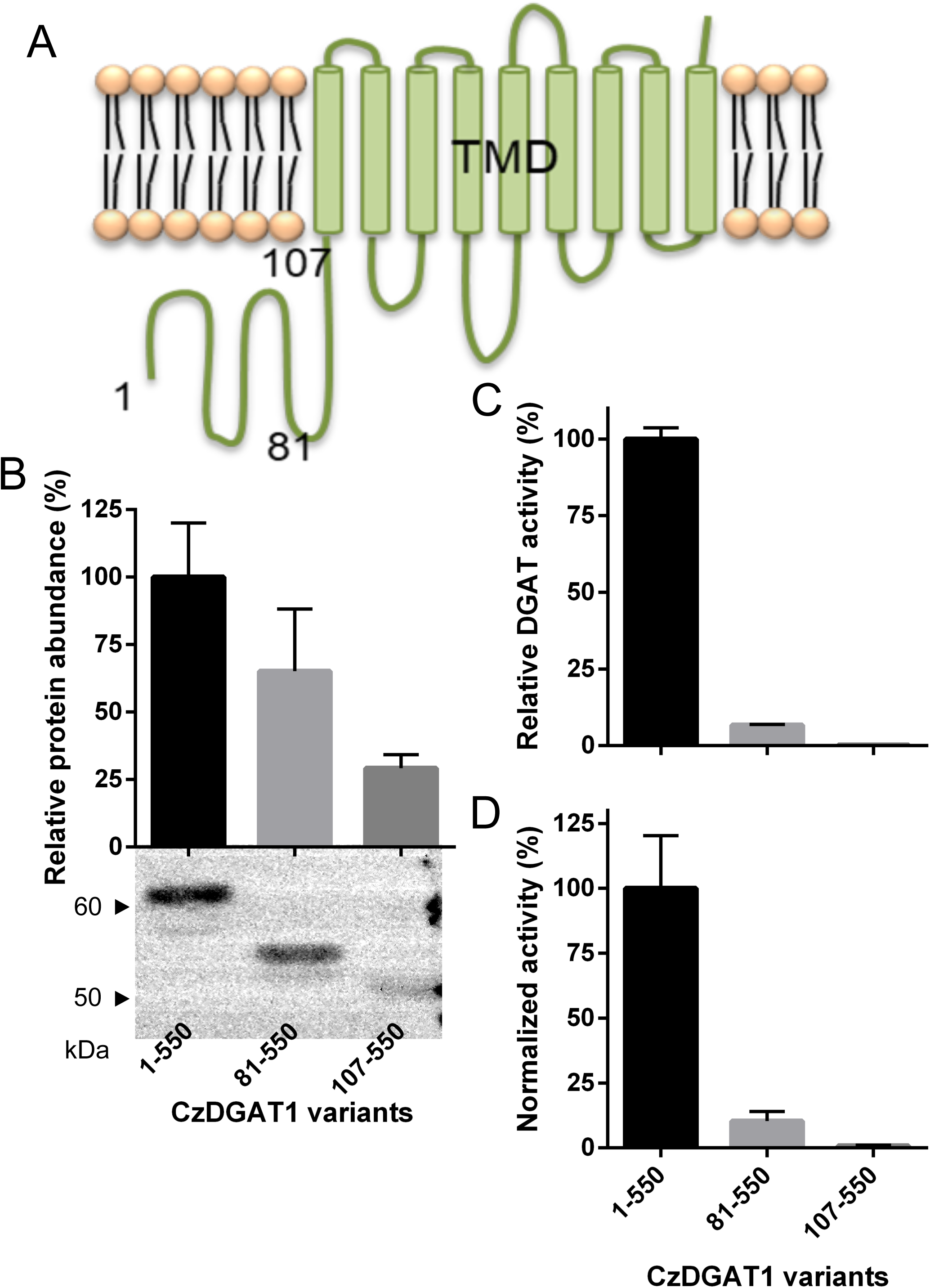
Truncation analysis of the N-terminal domain CzDGAT1. A, Predicted topology of CzDGAT1 by TMHMM (Krogh *et al*., 2001). CzDGAT1 has a 107-amino acid residue-long hydrophilic N-terminal region, followed by 9 predicted transmembrane domains (TMDs). The numbers indicate the different truncation points. B, Protein production level of the full-length (CzDGAT1_1-550_) and N-terminal truncated (CzDGAT1_81-550_ and CzDGAT1_107-550_) CzDGAT1 variant proteins. The enzyme amount was semi-quantified by Image J software (Schneider *et al*., 2012). C, *In vitro* DGAT activity of CzDGAT1_1-550_, CzDGAT1_81-550_ or CzDGAT1_107-550_. D, Normalized DGAT activity of CzDGAT1_1-550_, CzDGAT1_81-550_ or CzDGAT1_107-550_. Normalized activity was calculated by dividing the activity value by the corresponding protein accumulation. Data represent means ± S.D. (n = 3).

### Fusion with ACBP at the N-terminus of CzDGAT1 and its N-terminal truncation mutant enhanced the enzyme production

The acyl-CoA binding property of the N-terminal region has been revealed in *B. napus* DGAT1 (Weselake *et al*., 2006; Caldo *et al*., 2017). Considering the removal of the entire N-terminus of CzDGAT1 led to a nearly inactive enzyme, we further explored whether the N-terminal fusion of CzDGAT1_107-550_ with an ACBP would restore/improve the enzyme activity. We also tested if the fusion with ACBP could potentially improve DGAT performance by facilitating the channeling of acyl-CoA from the cytosol or the membrane lipid bilayer to the catalytic center of the enzyme. To test these hypotheses, AtACBP6, a 10-kDa soluble protein, was fused at the N-termini of CzDGAT1, CzDGAT1_81-550_ and CzDGAT1_107-550_, respectively, and the resulting fused proteins were produced in yeast H1246.

As shown in Figure 4A, N-terminal fusion with AtACBP6 largely increased the recombinant protein production in yeast, where the protein accumulation levels of AtACBP6 fused CzDGAT1, CzDGAT1_81-550_ and CzDGAT1_107-550_ were 5, 14, and 28-fold higher than the corresponding unfused CzDGAT1 variants, respectively. The specific activities of the AtACBP6 fused CzDGAT1 and CzDGAT1_81-550_ were also improved to about 2 and 5-fold greater than the unfused enzymes, respectively (Figure 4B). N-terminal fusion of CzDGAT1_107-550_ with AtACBP6, however, led to no improvement in enzyme activity compared to CzDGAT1_107-550_ (Figure 4B). The normalized enzyme activities of AtACBP6 fused CzDGAT1 and CzDGAT1_81-550_ were about 2.5 and 3-fold lower than the unfused enzymes, respectively (Figure 4C), suggesting that the enhanced enzyme activities of the fused proteins were mainly due to the enhanced protein abundance.

**Figure 4.**
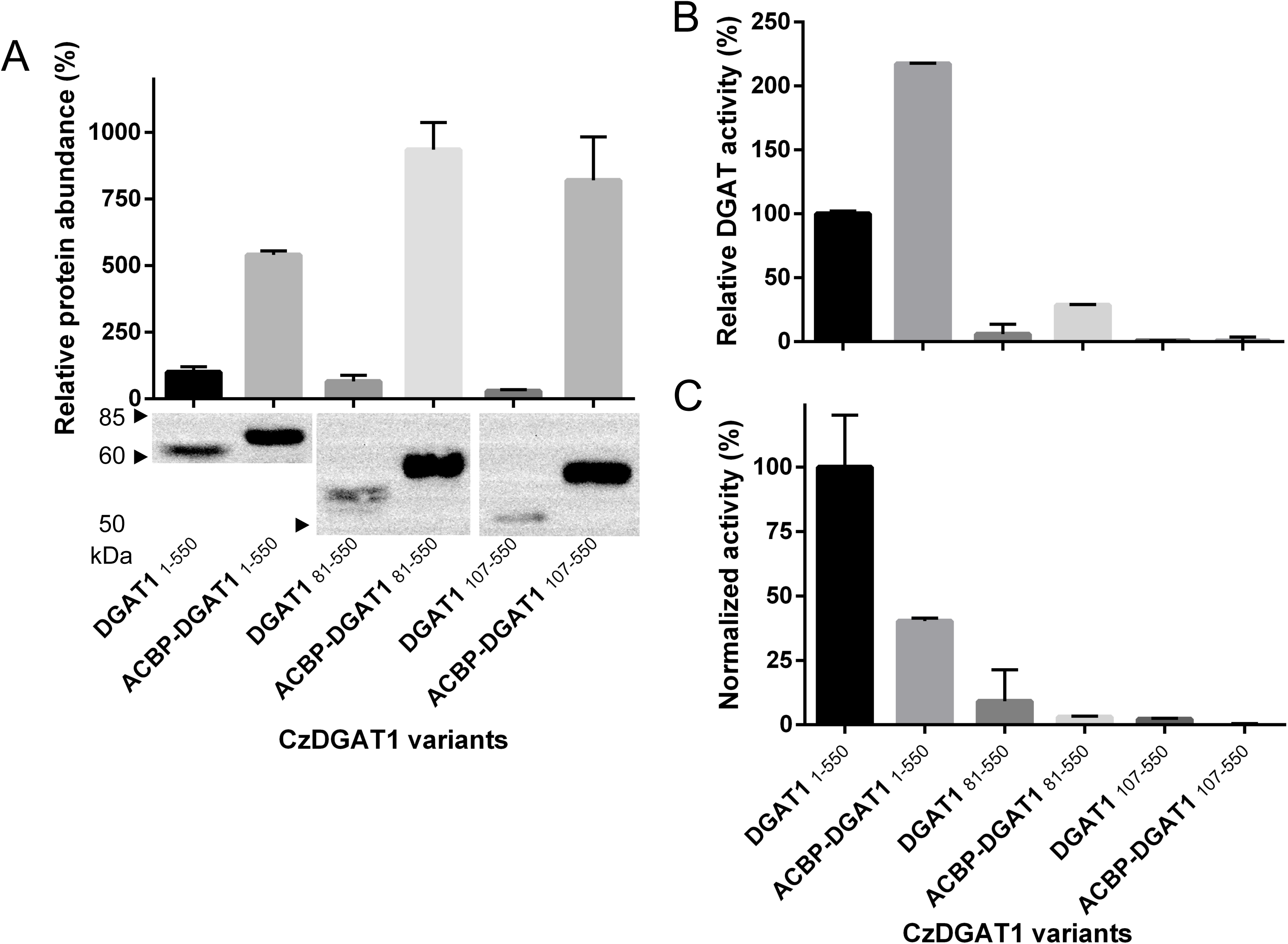
N-terminal fusion of an acyl-CoA binding protein (ACBP) with CzDGAT1 and its N-terminal truncation mutants. A, Protein production level of the full-length CzDGAT1 (DGAT1_1-550_), N-terminal truncated CzDGAT1 (DGAT1_81-550_ and DGAT1_107-550_) and their corresponding ACBP fused proteins (ACBP-DGAT1_1-550_, ACBP-DGAT1_81-550_ and ACBP-DGAT1_107-550_). The enzyme amount was semi-quantified by Image J software (Schneider *et al*., 2012). B, *In vitro* DGAT activity of the full-length, N-terminal truncated and ACBP fused CzDGAT1 proteins. C, Normalized DGAT activity of the full-length, N-terminal truncated and ACBP fused CzDGAT1 proteins. Normalized activity was calculated by dividing the activity value by the corresponding protein accumulation. Data represent means ± S.D. (n = 3).

### N-terminal fusion with ACBP kinetically improves CzDGAT1 and its N-terminal truncation mutant

To further explore the effects of fusing ACBP with DGAT1, the activities of the full-length CzDGAT1, ACBP-fused full-length CzDGAT1, CzDGAT1_81-550_, and ACBP-fused CzDGAT1_81-550_ were analyzed over increasing concentrations of oleoyl-CoA (Figures 5A-D). The full-length enzyme and the N-terminal truncation mutant had a similar response to the increasing acyl-donor concentration, with the maximum enzyme activity achieved at 5 and 7.5 *μ*M oleoyl-CoA, respectively. The ACBP-fused CzDGAT1_1-550_ and CzDGAT1_81-550_, on the other hand, reached the maximum enzyme activity at lower concentrations at 3 and 5 *μ*M oleoyl-CoA, respectively. The substrate saturation curves for CzDGAT1_1-550_ and CzDGAT1_81-550_ and their ACBP-fused versions had better fits to the allosteric sigmoidal equation over the Michaelis-Menten equation (Figures 5A-D). The Hill coefficients of CzDGAT1_1-550_, ACBP-fused CzDGAT1, CzDGAT1_81-550_ and ACBP-fused CzDGAT1_81-550_ were 1.39±0.06, 1.91±0.12, 1.71±0.06, and 1.57±0.14, respectively, which indicates that all these enzymes exhibited positive cooperativity. In addition, CzDGAT1_1-550_ and CzDGAT1_81-550_ had apparent S_0.5_ values of 1.48 ± 0.07 and 1.83 ± 0.05 *μ*M oleoyl-CoA, respectively (Table 1), suggesting that the oleoyl-CoA affinity of CzDGAT1_81-550_ might be lower than that of the full-length enzyme. Interestingly, ACBP-fused CzDGAT1_1-550_ and ACBP-fused CzDGAT1_81-550_ had apparent S_0.5_ values of 0.94 ± 0.04 and 1.50 ± 0.10 *μ*M oleoyl-CoA, respectively, which are 1.6 and 1.2-fold lower than the values of the corresponding unfused enzymes (Table 1). This result suggests that fusion with ACBP could enhance the affinity of DGAT variants to oleoyl-CoA.

**Figure 5.**
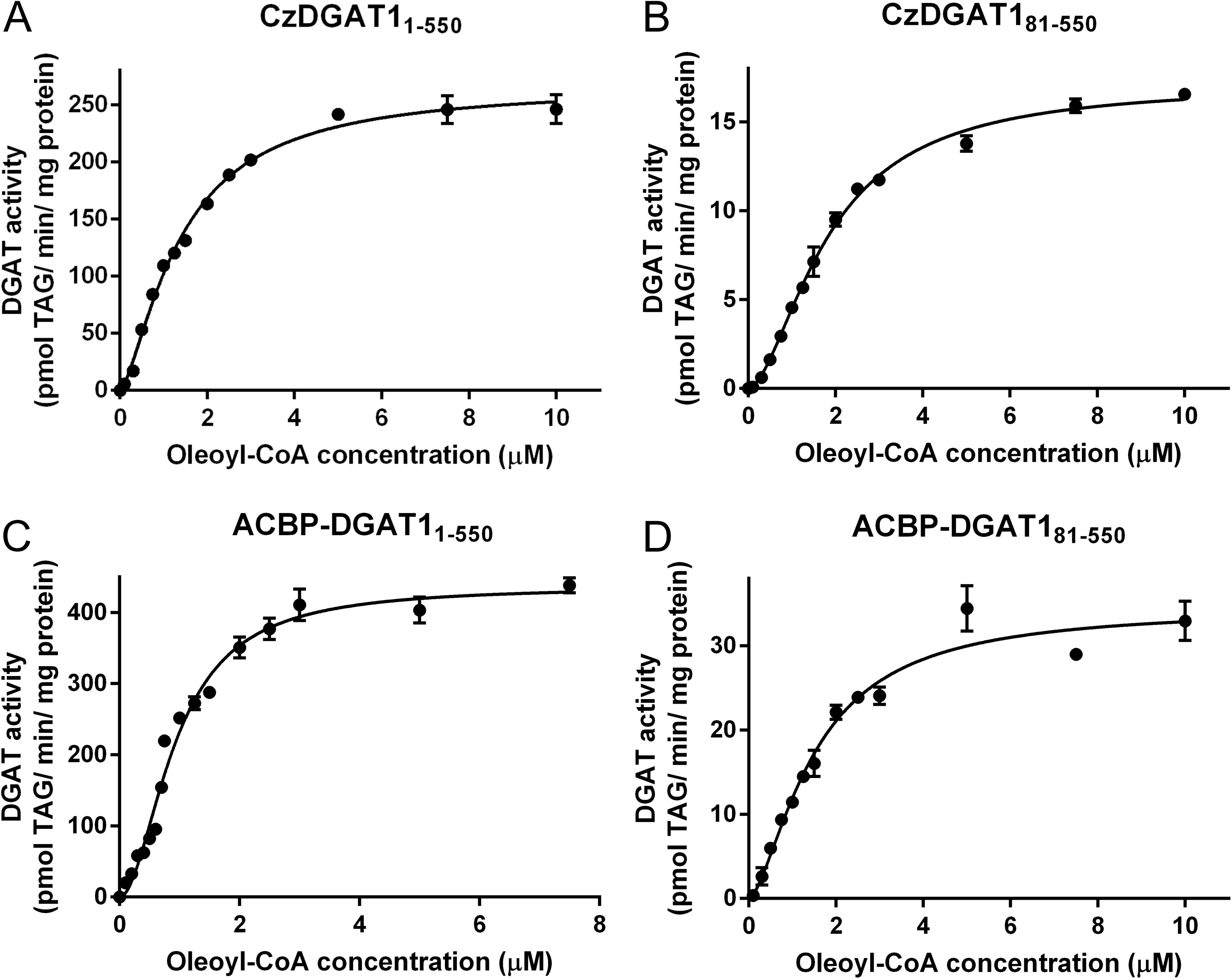
N-terminal fusion of an acyl-CoA binding protein (ACBP) with CzDGAT1 kinetically improves the enzyme. A-D, DGAT activities of the full-length CzDGAT1 (DGAT1_1-550_), N-terminal truncated CzDGAT1 (DGAT1_81-550_) and their corresponding ACBP fused proteins (ACBP-DGAT1_1-550_ and ACBP-DGAT1_81-550_) at oleoyl-CoA concentration from 0.1-7.5 or 10 µM. Data were fitted to the allosteric sigmoidal equation using GraphPad Prism. Data represent means ± S.D. (n = 3).

**Table 1.** Apparent kinetic parameters of CzDGAT1 variants. DGAT activity was examined at increasing oleoyl-CoA concentration from 0.1 to 7.5 or 10 μM. Data were fitted to a nonlinear regression using allosteric sigmoidal equation with GraphPad Prism software. Data shown are means ± S.D. (n=3).

| Enzyme | DGAT1<sub>1-550</sub> | DGAT1<sub>81-550</sub> | ACBP-DGAT1<sub>1-550</sub> | ACBP-DGAT1<sub>81-550</sub> |
| --- | --- | --- | --- | --- |
| Apparent $V_{\max}$ (pmol TAG/min/mg protein) | 276.8±6.94 | 17.14±0.28 | 436.7±10.1 | 34.54±1.27 |
| Hill coefficient | 1.39±0.06 | 1.71±0.06 | 1.91±0.12 | 1.57±0.14 |
| Apparent $S_{0.5}$ (μM) | 1.48±0.07 | 1.83±0.05 | 0.94±0.04 | 1.50±0.10 |
| Goodness of Fit/$R^2$ | 0.994 | 0.995 | 0.981 | 0.973 |

### N-terminal fusion with ACBP in CzDGAT1 boosts oil content in yeast and tobacco leaves

In order to determine whether the ACBP fused CzDGAT1 variants can boost oil production to higher levels compared to the native enzyme, cDNAs encoding the full-length CzDGAT1, the N-terminal truncated and the ACBP fused versions were individually introduced into the yeast strain H1246. Expression of *ACBP* or *LacZ* and co-expression of *CzDGAT1_1-550_* or *CzDGAT1_81-550_* with *ACBP* in yeast were used as controls. The production of CzDGAT1_1-550_ led to considerably increased neutral lipid accumulation in yeast at 24 h and 72 h, whereas no change in neutral lipid production was observed for the yeast expressing *CzDGAT1_81-550_* when compared to the LacZ control (Figure 6A). Fusion with ACBP at the N-terminal of CzDGAT1_1-550_ but not the N-terminal truncated enzyme was able to further increase the neutral lipid content, resulting in about 2-fold higher production than the native enzyme (Figure 6A). It should be noted that co-expression of *ACBP* with *CzDGAT1_1-550_* also promoted neutral lipid production in yeast to a comparable level to ACBP-fused CzDGAT1_1-550_ (Figure 6A), suggesting that ACBP may improve the acyl-CoA availability to DGAT and thus enhance the lipid production.

**Figure 6.**
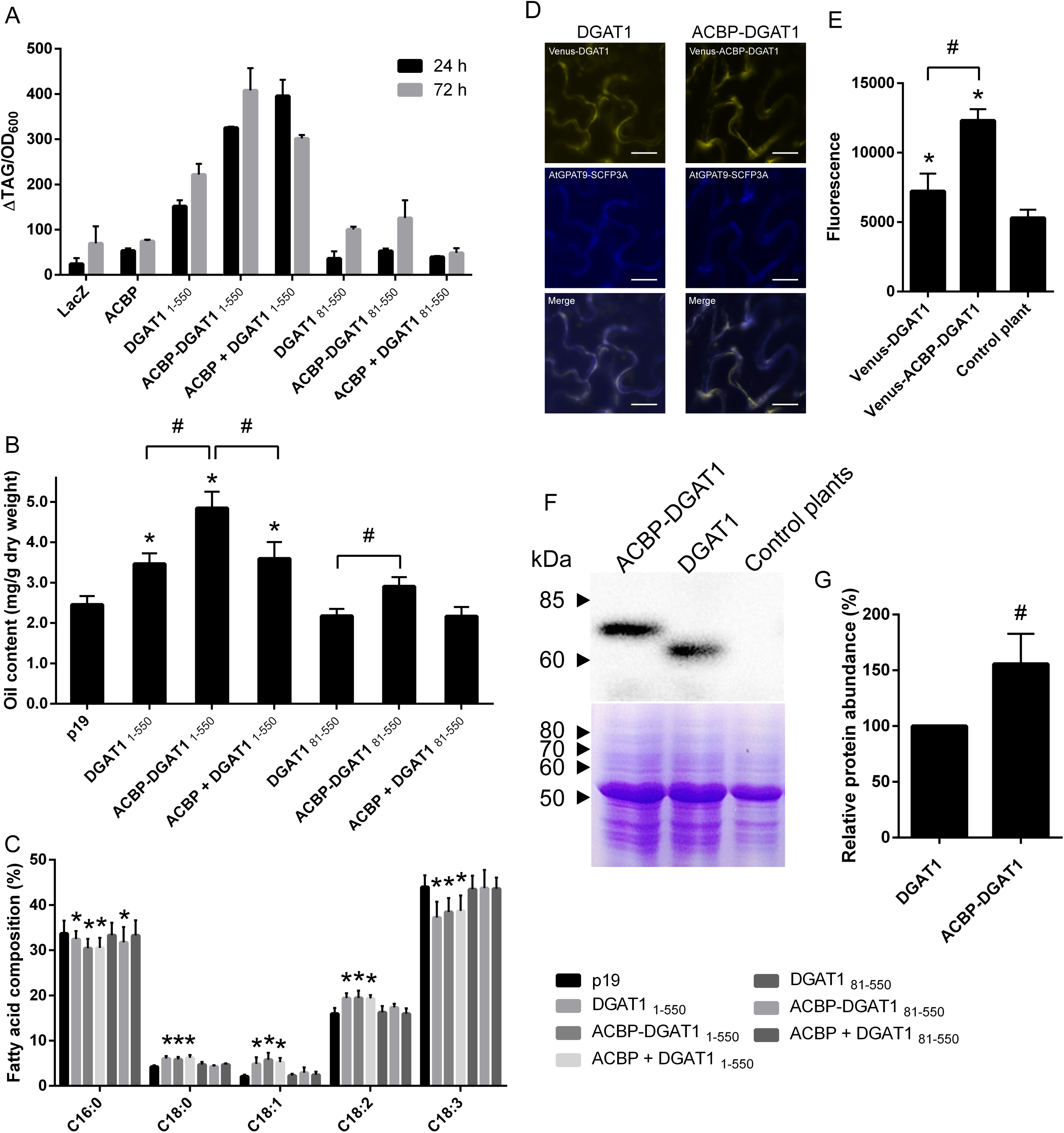
Fusion of acyl-CoA binding protein (ACBP) with CzDGAT1 boosts oil production in yeast and *Nicotiana benthamiana* leaves. A, Neutral lipid accumulation in yeast production ACBP-fused DGAT1. B, Triacylglycerol (TAG) content in transiently transformed *N. benthamiana* leaves. All cDNAs were constitutively expressed under the *CaMV 35S* promoter with the co-expression of the *p19* vector and *Arabidopsis thaliana Wrinkled1* (*AtWRI1*). C, Fatty acid composition of TAG in in transiently transformed *N. benthamiana* leaves. D, Subcellular localization of ACBP-fused DGAT1 in *N. benthamiana* leaf cells. Venus-tagged DGAT1 or ACBP-DGAT1 was co-localized with C-terminal SCFP3A tagged *Arabidopsis thaliana* glycerol-3-phosphate acyltransferase (AtGPAT9), a known endoplasmic reticulum (ER) localized protein. Scale bars represent 20 μm. E, Fluorescence intensity of ACBP-fused DGAT1. *N. benthamiana* leaves transiently expressing *Venus*-*DGAT1* or *Venus*-*ACBP*-*DGAT1* and the *p19* vector were used to quantify the fluorescent intensity. F, Western-blot immunodetection detection against V5 tagged ACBP-fused DGAT1 or DGAT1 on *N. benthamiana* leaf proteins. Coomassie Blue staining of duplicate leaf protein samples separated on SDS-PAGE gel is shown as a loading control. G, Relative protein accumulation levels of ACBP-fused DGAT1 or DGAT1 in *N. benthamiana* leaf cells based on western blot analysis. The enzyme amount was semi-quantified by Image J software (Schneider *et al*., 2012). C16:0, Palmitic acid; C18:0, Stearic acid; C18:1, Oleic acid; C18:2, Linoleic acid; C18:3, α-linolenic acid; p19, p19 + AtWRI1; DGAT1_1-550_, p19 + AtWRI1 + CzDGAT1_1-550_; ACBP-DGAT1_1-550_, p19 + AtWRI1 + ACBP fused CzDGAT1_1-550_; ACBP+DGAT1_1-550_, p19 + AtWRI1 + ACBP + CzDGAT1_1-550_; DGAT1_81-550_, p19 + AtWRI1 + CzDGAT1_81-550_; ACBP-DGAT1_81-550_, p19 + AtWRI1 + ACBP fused CzDGAT1_81-550_; ACBP+DGAT1_81-550_, p19 + AtWRI1 + ACBP + CzDGAT1_81-550_. Data represent means ± S.D. For A and G, n=3; for B and C, n=7; for E, n=4. The asterisk and pound sign indicate *P* < 0.05 as determined by paired one-tailed T-test.

To further examine the effects of CzDGAT1 variants on oil production in plant vegetative tissues, cDNAs encoding CzDGAT1_1-550_, ACBP-fused CzDGAT1_1-550_, CzDGAT1_81-550_ or ACBP-fused CzDGAT1_81-550_ were transiently co-expressed with *A. thaliana Wrinkled1* (*AtWRI1*) in the *N. benthamiana* leaves, respectively. *AtWRI1* encodes a transcription factor involved in the upregulation of genes in late glycolysis and fatty-acid biosynthesis (Xu, Caldo, *et al*., 2018) and co-expression of *AtWRI1* and *DGAT1* was previously found to substantially increase the TAG production in *N. benthamiana* leaves (Vanhercke *et al*., 2013). The over-expression of a cDNA encoding CzDGAT1_1-550_ or ACBP-fused CzDGAT1_1-550_ and the co-expression of *CzDGAT1_1-550_* and *ACBP* (ACBP + DGAT1_1-550_) led to considerable increases in leaf TAG content compared with the expression of *AtWRI1* alone (Figure 6B). Interestingly, ACBP-fused CzDGAT1_1-550_ had a higher impact on improving the leaf TAG production than CzDGAT1_1-550_ or the co-expression group (ACBP + DGAT1_1-550_), where a 1.40 or 1.35-fold increase in TAG content was observed, respectively (Figure 6B). Consistently, the TAG fatty acid composition was also affected in the groups with increased TAG content compared with the *AtWRI1* control group, in which palmitic acid (C16:0) and C18:3 contents were decreased while the content of stearic acid (C18:0), oleic acid (C18:1*Δ^cis9^*, C18:1), and C18:2 were increased (Figure 6C). The over-expression of a cDNA encoding ACBP-fused CzDGAT1_81-550_ also resulted in a slight increase in TAG production relative to that of the unfused enzyme, although both groups showed no significant difference in TAG content and composition compared to the *AtWRI1* control group, with an exception of a decrease in C16:0 content in the ACBP-fused CzDGAT1_81-550_ (Figures 6B and C).

To test whether ACBP fusion affects the subcellular localization of CzDGAT1, Venus, a variant of yellow fluorescent protein, was fused to the N-terminal of CzDGAT1_1-550_ or ACBP-CzDGAT1_1-550_, and was transiently co-produced with *A. thaliana* glycerol-3-phosphate acyltransferase (AtGPAT9) containing a C-terminal SCFP3A (a cyan fluorescent protein) fusion in tobacco leaves, which is known to reside in the endoplasmic reticulum (ER) (Gidda *et al*., 2009). Both CzDGAT1_1-550_ or ACBP-CzDGAT1_1-550_ were found to co-localize with AtGPAT9 (Figure 6D), suggesting their ER localization. It should also be noted that linkage of ACBP to the N-terminus of CzDGAT1 enhanced the CzDGAT1 production in tobacco leaves as indicated by the increased fluorescence intensity (Figure 6E) and enhanced protein accumulation levels based on Western blot analysis (Figures 6F and G).

## DISCUSSION

Recently, DGAT1s from a growing number of microalgal species have been characterized with the focus on elucidating their physiological role and application in manipulating oil production (Kirchner *et al*., 2016; Wei *et al*., 2017; Guo *et al*., 2017; Guihéneuf *et al*., 2011). The structure-function perspectives of algal DGAT1 and the relationship to plant and animal DGAT1, however, remain largely unexplored. In the current study, the evolutionary and structural features of algal DGAT1 in comparison to the higher eukaryotic enzymes were investigated via *in silico* analysis, truncation mutagenesis and *in vitro* enzyme assay, using DGAT1 of the emerging model green microalga *C. zofingiensis* (Roth *et al*., 2017; Mao *et al*., 2019; Liu *et al*., 2019) as a representative. Moreover, an improved DGAT1 variant was engineered by N-terminal fusion with ACBP, and its potential in enhancing oil production was explored using yeast expression and *N. benthamiana* transient expression systems.

The evolutionary and sequence analyses revealed that CzDGAT1 and other green microalgal DGAT1 are closely related to the plant DGAT1 clade. In addition, algal DGAT1 has similar structural features to plant and animal DGAT1, such as the highly conserved C-terminal hydrophobic region forming multiple TMDs and the variable hydrophilic N-terminal region (Figures 1 and S1 and Table S1). Interestingly, the N-terminus of algal DGAT1 varies dramatically in length ranging from 20 to 326 amino acid residues, which differs from that of plant and animal DGAT1 (Figure 1 and Table S1). The large variation in the N-terminal size appears to be not relevant to the maintenance of enzyme activity, since many DGAT1s with extremely long or short N-termini have been demonstrated to function in TAG biosynthesis (Kirchner *et al*., 2016; Wei *et al*., 2017; Mao *et al*., 2019). Furthermore, the N-terminal region of algal DGAT1 is likely to have different structural features from that of plant and animal DGAT1. The majority of the N-terminus of plant and animal DGAT1 is disordered, while the small folded portion contains a conserved allosteric site (Caldo *et al*., 2017). On the contrary, the algal DGAT1 N-terminal region has much less propensity to be disordered based on secondary structure analysis (Figures 2E, F and S2) and the possible allosteric site is less conserved in algal DGAT1 (Figure 2D). Taken together, these results suggest that the N-terminal region of algal DGAT1 may have evolved differently from that of plant and animal in terms of structure and function.

The distinct features of the N-terminal domain of algal DGAT1 in enzyme catalysis are also supported by the evidence from the truncation mutagenesis of CzDGAT1. Previous study on *B. napus* DGAT1 suggested that the first 80 amino acid residues in the N-terminal region down regulates enzyme activity and the removal of that region led to higher enzyme activity (Caldo *et al*., 2017). CzDGAT1 has a N-terminal region (1-107) of a similar size to that of plant and animal DGAT1 (Table S1), but truncation of the equivalent region to the autoinhibitory motif in *B. napus* DGAT1 (CzDGAT1_81-550_; Figure 2D) diminished the enzyme activity by 10-fold (Figure 3D), suggesting that this region may not function as an autoinhibitory motif in CzDGAT1. Indeed, the DISOPRED analysis suggested that the N-terminal domain of CzDGAT1 does not have a strong tendency to have the disordered region where autoinhibitory motifs are normally found (Figures 2E and F). Moreover, the N-terminal domain was found to be involved in mediating positive cooperativity in *B. napus* DGAT1, where the N-terminal truncated *B. napus* DGAT1_81-501_ had a decreased Hill coefficient value compared to the full-length enzyme (Caldo *et al*., 2017). This appears not to be the case for CzDGAT1 since both CzDGAT1 and CzDGAT1_81-550_ exhibited positive cooperativity with close Hill coefficient values (Figures 5A and B, Table 1). Interestingly, N-terminal truncation of CzDGAT1 (CzDGAT1_81-550_) seems to alleviate the substrate inhibition of the full-length enzyme (Figure S3 and Table S2), suggesting that segment 1-80 may be related to a low affinity non-catalytic acyl-CoA binding site. Furthermore, the N-terminal domains of *B. napus* and mouse DGAT1 were shown to be responsible for oligomer formation (McFie *et al*., 2010; Weselake *et al*., 2006; Caldo *et al*., 2017). Although whether this domain functions in self-association in CzDGAT1 remains further exploration, DISOPRED analysis showed that the N-terminus of CzDGAT1 has a less propensity to form protein-protein interactions than *B. napus* and mouse DGAT1 (Figures 2E, F and S2). In addition, truncation of this domain did not affect the self-interaction of CzDGAT1 in the membrane yeast two-hybrid assay (Figure S4). It is also possible that the observed self-interaction was due to the presence of a region in the hydrophobic part of CzDGAT1 that is involved in oligomer formation. Indeed, a 16-kDa fragment of mouse DGAT1 present in the ER lumen was also found to likely form dimer/tetramer (McFie *et al*., 2010).

The removal of the entire N-terminal region in CzDGAT1 (CzDGAT1_107-550_) almost inactivated the enzyme (Figure 3D), suggesting that the N-terminal region is not necessary for enzyme catalysis but is important for maintaining high enzyme activity. This agrees with the previous studies on the *B. napus* DGAT1 (Caldo *et al*., 2017). Previously, the N-termini of *B. napus* and mouse DGAT1 were found to associate with acyl-CoA in a sigmoidal fashion (Weselake *et al*., 2006; Siloto *et al*., 2008). Replacement of the N-terminal region by ACBP in CzDGAT1 (ACBP fused CzDGAT1_107-550_), however, was not able to restore or improve the enzyme activity (Figure 4), suggesting that the N-terminus may function as a regulatory domain rather than as an acyl-CoA binding site. Indeed, the last 20-30 amino acid residues in the N-terminus of *B. napus* DGAT1 has a well-folded structure and constitutes a regulatory domain for allosteric binding of acyl-CoA and CoA, where the binding of CoA would trigger the subsequent inhibition of DGAT activity (Caldo *et al*., 2017). The allosteric site for CoA, however, is only partially conserved in CzDGAT1 (Figure 2D), and the activity of CzDGAT1 was not affected in the presence of 50 µM CoA (Figure S5), suggesting that the allosteric site for CoA might be not present in the algal DGAT1 or not activated under this experimental condition. The CoA inhibition was also not triggered by fusion with ACBP to CzDGAT1 (Figure S5), despite that ACBP is able to bind CoA alone (Robinson *et al*., 1996). However, we cannot rule out the possibility that failure in restoring DGAT activity by replacing the N-terminus with ACBP may be caused by the high binding affinity of ACBP to acyl-CoAs, while a moderate binding affinity being required for the enzyme function. A comprehensive characterization of the N-termini of CzDGAT1 and other algal DGAT1s with respective to the acyl-CoA binding ability would be an interesting next step to explore the possible role of this domain in the catalysis of algal DGATs.

The two isoforms of DGAT1 from the green microalga *C. zofingiensis* (Mao *et al*., 2019) have very distinct features. They share 39.9% amino acid pairwise identity and have very different lengths mainly attributable to the variable N-terminal regions (Figure S1). The N-terminus of the CzDGAT1 isoform studied in the current study is of a similar length to that of plant and animal DGAT1 but with less tendency to be disordered, whereas the other isoform has an extremely long hydrophilic N-terminus of 289 amino acid residues with the first 160 amino acid residues showing low tendency to become disordered and the segment consisting of the remaining 129 residues being highly disordered (Table S1 and Figure S2). Since DGAT1 isoform with a longer N-terminus was found to have a higher ability to restore TAG biosynthesis in the yeast mutant (Mao *et al*., 2019), it is possible that their different N-terminal domain features may directly impact on the enzyme activity and accumulation. Therefore, it would be interesting to comprehensively compare the structure and function features of the N-terminal domains of both CzDGAT1 isoforms in the future. Additionally, it may also be worthwhile to explore the activities and physiological functions of CzDGAT1 in detail, especially considering that the coding genes of both isoforms were up-regulated under nitrogen deprivation and high-light stress conditions to contribute to TAG accumulation but with very different transcript abundances (under both conditions) and response patterns (under high-light stress) (Roth *et al*., 2017; Mao *et al*., 2019; Liu *et al*., 2019).

It is interesting to note that N-terminal fusion with ACBP could markedly increase the yeast recombinant protein production up to 28 folds (Figure 4A). It has been suggested that the identity of the amino acid residues at the N-terminus of proteins could potentially affect the protein turnover and/or translation rate and thus enhance the protein production (Greer *et al*., 2015; Sriram *et al*., 2011). Indeed, the addition of N-terminal tag was found to increase the production of *B. napus* DGAT1 and plant fatty acid desaturases in yeast cells (O’Quin *et al*., 2009; Greer *et al*., 2015). Previously, fusion with small soluble proteins, such as thioredoxin, ubiquitin and maltose binding protein was also found to augment the accumulation levels of recombinant protein in *Escherichia coli* and/or yeast likely by improving protein stability and/or translation efficiency (Marsh *et al*., 1989; Pryor and Leiting, 1997; Jacquet *et al*., 1999; Ecker *et al*., 1989). The protein fusion strategy, especially fusion to ubiquitin, has also met considerable success in increasing protein production in transgenic plants (Mishra *et al*., 2006; Tian and Sun, 2011; Hondred *et al*., 1999; Streatfield, 2007). Our finding that linkage to ACBP enhanced the CzDGAT1 production in tobacco leaves (Figures 6E-G) suggests that ACBP might also have potential to be used as a protein fusion partner to enhance protein accumulation in plants, especially for membrane-bound acyl-CoA-dependent enzymes.

Fusion with ACBP not only augmented the production of CzDGAT1 (Figures 4A and 6E-G) but also improved the kinetic parameters of the enzyme (Figure 5). Our kinetic analysis showed that ACBP-fused CzDGAT1_1-550_ or ACBP-fused CzDGAT1_81-550_ still exhibited positive cooperativity despite that ACBP binds acyl-CoA in a typical hyperbolic manner (Brown *et al*., 1998; Yurchenko *et al*., 2009) and, more importantly, the fused enzymes had increased affinity for oleoyl-CoA with the apparent S_0.5_ values decreasing by 1.6 and 1.2-fold, respectively (Table 1 and Figure 5). DGAT affinity for oleoyl-CoA has been reported as an important determinant of oil production since a strong correlation of oleoyl-CoA affinity with oil content was found in soybean expressing *DGAT1* variants (Roesler *et al*., 2016). Cytosolic 10-kDa ACBP, consisting of four α helixes (Figure S6), is capable of binding acyl-CoAs with high affinity (Robinson *et al*., 1996; Du *et al*., 2016). Considering the increased DGAT affinity for oleoyl-CoA of the ACBP-fused CzDGAT1, we proposed that ACBP would facilitate the feeding of acyl-CoA to the catalytic pocket of CzDGAT1 via capturing cytosolic acyl-CoAs (or those partitioned into the membrane lipid bilayer) and subsequent channeling to DGAT by proximity (Figure S6). Similarly, fusion of proteins in a consecutive reaction has been shown to have synergistic effects in substrate conversion in a single catalytic region (Elleuche, 2015). ACBP has been implicated in acyl-CoA binding and transport, which maintains the substrate supply for the acyl-CoA-dependent acyltransferases in the ER, including DGAT (Yurchenko and Weselake, 2011). Recently, ACBP2 was found to probably interact directly with lysophospholipase 2 in *A. thaliana* and thereby facilitate the lysophosphatidylcholine hydrolysis (Miao *et al*., 2019). Therefore, it is plausible to assume the presence of transient interactions between ACBP and DGAT, the enhancement of which via protein fusion may further lead to an efficient substrate channeling. Furthermore, *B. napus* ACBP has been shown to slightly stimulate DGAT1 activity when the concentration of acyl-CoA is higher than that of ACBP, but to inhibit DGAT activity when ACBP is in excess likely due to the competition of ACBP with ACBP-bound acyl-CoA or DGAT for enzyme-substrate interaction (Yurchenko and Weselake, 2011). ACBP-fused CzDGAT1 or CzDGAT1_81-550_, on the other hand, showed a more rapid response to increasing acyl-CoA concentration from 0.1 to 5 or 10 µM than the unfused enzymes, respectively (Figure 5 and Table 1), suggesting that in the 1:1 fusion form ACBP may facilitate enzyme-substrate binding rather than compete with DGAT for substrate.

The kinetically improved DGAT1 via fusion with ACBP shows promises in oil production in both yeast and plant systems (Figure 6). Although ACBP-fused DGAT1 was able to augment the yeast neutral lipid production up to two-fold relative to that of the unfused enzyme, a comparable level was also achieved in the yeast co-expressing individual fusion groups (Figure 6A). In tobacco transient expression system, ACBP-fused DGAT1 was more efficient to boost leaf TAG content than the unfused enzyme and the co-expressed group. It should be noted that the sizes of the acyl-CoA pool in yeast and tobacco leaf cells are different. The cellular concentration of total acyl-CoA pool in *S. cerevisiae* has been reported to be in the range of 10-42 µM depending on the strain and its metabolic state (Schjerling *et al*., 1996; Mandrup *et al*., 1993; Knudsen *et al*., 1994); whereas the concentration of total acyl-CoAs in tobacco leaves was determined to be 0.5 µM (Moreno *et al*., 2014) [intracellular concentrations were converted to µM assuming a specific cell volume of 3.7 × 10^-14^ L/cell (Krink-Koutsoubelis *et al*., 2018) and 1 mg equal to 1 μL volume (Larson and Graham, 2001), for yeast and plant cells, respectively]. Co-production of ACBP and DGAT1 appears to work better in the cells with the acyl-CoA concentration at higher levels rather than lower levels (Figures 6A and B). It is possible that co-production of ACBP with DGAT1 affected *in vivo* DGAT1 activity in a manner dependent on the cellular acyl-CoA concentration, where co-produced ACBP stimulates DGAT1 activity at high acyl-CoA concentration but restricts DGAT1 activity at low acyl-CoA concentration, which agrees with the previous finding on the *in vitro* stimulation of DGAT activity by *B. napus* ACBP (Yurchenko and Weselake, 2011). ACBP-fused DGAT1, on the other hand, showed good performance in cells with both low and high concentrations of acyl-CoAs (Figures 6A and B) likely due to the improved enzyme kinetics and protein production levels, despite that fusion with ACBP slightly enhanced the substrate inhibition of DGAT1 at the acyl-CoA concentrations above 10 µM (Figure S3 and Table S2). Moreover, it would be interesting to explore the potential of co-producing DGAT with other ACBP isoforms, such as ER-bound ACBP, in improving TAG production, since both soluble and ER-bound ACBP may be involved in the capture and shuttle of acyl-CoAs for downstream acyl-CoA-dependent acyltransferases.

In conclusion, our results suggested that algal DGAT1 may have different evolutionary and structural features from plant and animal DGAT1 with respect to the hydrophilic N-terminal domain. This domain is predicted to be present in a less disordered state in CzDGAT1 than that of animal/plant DGAT1. Although the N-terminal domain is not necessary for acyltransferase activity of CzDGAT1, its removal, however, led to huge decreases in CzDGAT1 enzyme activity, which cannot be restored by fusion with an ACBP. We also found that fusion of ACBP to the N-terminus of the full-length CzDGAT1 could not only augment the protein accumulation levels in yeast and tobacco leaves but also kinetically improve the enzyme. ACBP-fused DGAT1 was more effective in improving the oil contents of yeast cells and vegetative tissues than the native DGAT1. This strategy may have great potential in engineering membrane-bound acyl-CoA-dependent enzymes and manipulating oil biosynthesis in plants, algae and other oleaginous organisms.

## EXPERIMENTAL PROCEDURES

### Sequence analysis

Multiple sequence alignments of DGAT1 proteins from different animal, plant and microalgal species (Table S1) were performed using the L-INS-i method implemented in MAFFT v7.271 (Katoh and Standley, 2013) and trimmed using trimAl v1.2 (Capella-Gutiérrez *et al*., 2009). The resulting alignments were used for model selection using IQ-TREE (v1.3.11.1) with the option “- m TESTONLY”. The phylogenetic tree was then constructed with the best-fit model for protein alignment (LG+F+I+G) using Phyml v3.0 (Guindon *et al*., 2010) and visualized with iTOL v3 (Letunic and Bork, 2016). The topology organization of DGAT1 was predicted using TMHMM (Krogh *et al*., 2001).

### Construct preparation, yeast transformation and heterologous expression of *DGAT1* variants

The coding sequence of *CzDGAT1* (Phytozome accession number: Cz09g08290) was chemically synthesized (General Biosystems, Morrisville, NC), and re-cloned into the pYES2.1 yeast expression vector (Invitrogen, Burlington, Canada) under the control of the galactose-inducible *GAL1* promoter. N-terminal truncation mutants of *CzDGAT1* were PCR-amplified and subcloned into the pYES2.1 vector. The coding sequence of *AtACBP6* (NCBI accession number: NM_102916) was PCR-amplified from the plasmid pAT332 (kindly provided by Dr. Mee-Len Chye, University of Hong Kong) (Chen *et al*., 2008), and subcloned into the pYES2.1 vector and the pESC-leu2d(empty) vector (a gift from Dr. Jay Keasling; Addgene plasmid # 20120) (Ro *et al*., 2006). To generate ACBP-CzDGAT1_1-550_, ACBP-CzDGAT1_81-550_ and ACBP-CzDGAT1_107-550_ fusion proteins, the coding sequences of *AtACBP6* and variant *CzDGAT1s* were individually amplified and the resulting amplicons were fused using overlap extension PCR. The DNA sequences of ACBP-CzDGAT1_1-550_, ACBP-CzDGAT1_81-550_ and ACBP-CzDGAT1_107-550_ fusion proteins were then subcloned into the pYES2.1 vector. The stop codon was removed from each sequence for in-frame fusion with a C-terminal V5 tag, which is encoded in the pYES2.1 vector. The primers used for the preparation of all the constructs are listed in Table S3.

After the integrity of each construct was confirmed by sequencing, the constructs were transformed into the quadruple mutant strain *S. cerevisiae* H1246 (*MATα are1-Δ::HIS3, are2-Δ::LEU2, dga1-Δ::KanMX4, lro1-Δ::TRP1 ADE2*) using an *S.c.* EasyComp Transformation Kit (Invitrogen) for yeast heterologous expression (Xu, Holic, *et al*., 2018; Xu *et al*., 2017; Xu *et al*., 2019). The recombinant yeast cells were first cultured overnight in liquid minimal medium containing 0.67% (w/v) yeast nitrogen base, 0.2% (w/v) synthetic complete medium lacking uracil (SC-Ura) and 2% (w/v) raffinose. The yeast cultures were then used to inoculate the induction medium containing 0.67% (w/v) yeast nitrogen base, 0.2% (w/v) SC-Ura, 2% (w/v) galactose, and 1% (w/ v) raffinose at an initial optical density of 0.4 at 600 nm (OD_600_). For co-expression of *AtACBP6* and *CzDGAT1*, SC-Ura in the liquid minimal medium was replaced by synthetic complete medium lacking leucine and uracil. For the fatty acid feeding experiment, yeast cells were cultured in the induction medium with the supplementation of 200 mM of C18:2 or C18:3 fatty acid. Yeast cultures were grown at 30°C with shaking at 220 rpm.

### Preparation of yeast microsomal fractions

Microsomal fractions containing recombinant CzDGAT1 variants were isolated from yeast cells as described previously (Xu *et al*., 2017; Xu *et al*., 2019; Xu, Holic, *et al*., 2018). In brief, the recombinant yeast cells were collected at the mid log growth stage with an OD600 value around 6.5, washed, and then resuspended in 1 mL of lysis buffer [20 mM Tris-HCl pH 7.9, containing 10 mM MgCl_2_, 1 mM EDTA, 5% (v/v) glycerol, 300 mM ammonium sulfate and 2 mM dithiothreitol]. The cells were homogenized by a bead beater (Biospec, Bartlesville, OK, USA) in the presence of 0.5 mm glass beads and then centrifuged at 10 000 g at 4°C for 30 min to remove cell debris and glass beads. The recovered supernatant was further centrifuged at 105 000 g at 4°C for 70 min to pellet the microsomes. The resulting microsomal fractions were resuspended in ice-cold suspension buffer (3 mM imidazole buffer, pH 7.4, and 125 mM sucrose). Protein concentration was quantified by the Bradford assay (Bio-Rad, Mississauga, Canada) using BSA as a standard (Bradford, 1976).

### *In vitro* DGAT1 assay

*In vitro* DGAT assay was performed according to the procedure described previously (Xu *et al*., 2017; Xu *et al*., 2019; Xu, Holic, *et al*., 2018). Briefly, a 60-*µ*L reaction mixture containing 200 mM HEPES-NaOH (pH 7.4), 3.2 mM MgCl_2_, 333 *µ*M *sn*-1,2-diolein [dispersed in 0.2% (v/v) Tween 20], 15 *µ*M [1-^14^C] oleoyl-CoA (56 *μ*Ci/*μ*mol; American Radiolabeled Chemicals, St. Louis, MO, USA), and 10-50 *µ*g of microsomal protein was incubated at 30°C for 4-30 min with shaking. The reaction was initiated by adding microsomes containing recombinant CzDGAT1 variants and terminated by adding 10 *µL* of 10% (w/v) SDS. The entire reaction mixture was then loaded onto a thin-layer chromatography (TLC) plate (0.25 mm Silica gel, DC-Fertigplatten, Macherey-Nagel, Germany). The plate was developed with hexane/diethyl ether/acetic acid (80:20:1, v/v/v) and the resolved lipids were visualized by phosphorimaging (Typhoon Trio Variable Mode Imager, GE Healthcare, Mississauga, Canada). The corresponding TAG spots were scraped and quantified for radioactivity using a LS 6500 multi-purpose scintillation counter (Beckman-Coulter, Mississauga, Canada).

For kinetic assay, the concentration of [1-^14^C] oleoyl-CoA was varied from 0.1 to 25 *µ*M while *sn*-1,2-diolein was held constant at 333 *µ*M. The DGAT assay reaction time and the quantity of microsomal protein used were as follows: for CzDGAT1_1-550_ and ACBP-CzDGAT1_1-550_, 10 *µ*g of microsomal protein for 4 min; for ACBP-CzDGAT1_81-550_, 5 *µ*g of microsomal protein for 30 min; for CzDGAT1_81-550_, 10 *µ*g of microsomal protein for 30 min. Enzyme kinetic parameters were calculated by fitting the data to the Michaelis-Menten equation, allosteric sigmoidal equation, or a previously proposed model accounting for sigmoidicity and substrate inhibition (Xu *et al*., 2017) using the program GraphPad Prism (version 6.0; GraphPad Software, La Jolla, CA, USA).

Transient expression of *CzDGAT1* variants in *N. benthamiana* leaves *N. benthamiana* plants were grown in a growth chamber at 25°C, 50% humidity and 16/8 hr day/night cycle. For transient expression in *N. benthamiana* leaves for lipid production, cDNAs encoding *AtWRI1* (previously isolated using *A. thaliana* cDNA in our lab), CzDGAT1_1-550_, the N-terminal truncation mutant (CzDGAT1_81-550_) and their ACBP-fused versions (ACBP-CzDGAT1_1-550_ and ACBP-CzDGAT1_81-550_) were subcloned in a pGREEN 0229 vector under a *cauliflower mosaic virus* (*CaMV*) *35S* promoter, respectively. For the examination of the subcellular localization, cDNAs encoding CzDGAT1_1-550_ and ACBP-CzDGAT1_1-550_ were fused in frame to the downstream of *Venus*, which was amplified from the pSYFP2-SCFP3A plasmid (a gift from Dr. Dorus Gadella; Addgene plasmid # 22905) (Kremers *et al*., 2006), and was inserted downstream of a *CaMV 35S* promoter in the modified pGPTVII vector (kindly provided by Dr. Jörg Kudla, University of Münster) (Becker *et al*., 1992; Gehl *et al*., 2009). The coding sequence of *AtGPAT9* (Singer *et al*., 2016) was fused in frame to the upstream of *SCFP3A* (amplified from the pSYFP2-SCFP3A plasmid), subcloned in the modified pGPTVII vector under a *CaMV 35S* promoter and used as an ER marker (Gidda *et al*., 2009).

All constructs were individually transformed to *Agrobacterium tumefaciens* GV3101 cells via electroporation. Each pGREEN construct was transformed along with the pSOUP helper plasmid. *A. tumefaciens* cultures containing the *p19* vector encoding a viral suppressor protein and each variant *CzDGAT1* were mixed in a transformation medium [50 mM MES, 2 mM Na_3_PO_4_, 0.5% (w/v) glucose and 0.1 mM acetosyringone] with the final OD_600_ of each culture equal to 0.125 (or 0.25 for subcellular localization experiments) prior to infiltration into *N. benthamiana* leaves as described by Vanhercke et al. (2013). For lipid analysis, *N. benthamiana* plants were grown for a further five days before leaf samples were collected, flash frozen, freeze-dried, and stored at −80°C. For protein extraction, *N. benthamiana* leaves were harvested after 2 days of infiltration and were ground in liquid nitrogen to a fine powder using mortar and pestle. The resulting leaf powders were then mixed with 1 volume (v/w) of ice-cold extraction buffer (4 M urea, 100 mM DTT, 1% Triton X-100, and 1 mM PMSF), and incubated on ice for 10 min. The homogenate was clarified by centrifugation at 12 000 g for 5 min at 4°C and the supernatant was used for SDS-PAGE gel analysis immediately after preparation.

For subcellular localization experiments, *AtGPAT9*-*SCFP3C* was co-infiltrated with the *p19* vector and *Venus*-*CzDGAT1* or *Venus-ACBP-CzDGAT1* and the fluorescence of the lower epidermis of leaves after 2-3 days of infiltration was visualized using a fluorescent microscope (Axio Imager M1m microscope; Carl Zeiss Inc., Germany). The excitation wavelengths for Venus and SCFP3A were 546/12 and 365 nm, respectively, and the emission filter wavelengths were 575–640 nm for Venus and 455/50 nm for SCFP3A. For the quantification of fluorescence intensity, the fluorescent proteins were extracted from tobacco leaves transiently expressing the *p19* vector and *Venus*-*CzDGAT1* or *Venus*-*ACBP-CzDGAT1* by grinding on ice with a two-fold volume of pre-chilled extraction buffer containing 100 mM MOPS (pH 7.2), 5 mM MgCl_2_, 0.02% BSA and 1% protease inhibitor, followed by centrifugation. One hundred microliters of the supernatant were then placed into 96-well solid black plates (Corning Inc., Corning, NY, USA) and the fluorescence intensity of Venus-tagged protein was measured on a Synergy H4 Hybrid reader (Biotek, Winooskit, VT, USA) at excitation and emission wavelengths of 485 nm and 528 nm, respectively.

### Western blotting

Equivalent amounts of yeast microsomal proteins (15 *µ*g) or protein extracts from leaf samples containing recombinant CzDGAT1 variants were incubated with 5× SDS loading buffer at room temperature for 15 min, resolved through SDS-PAGE Gels (Bio-Rad) and electrotransfered (2 h at 80 V or 16 h at 30 V and 4°C) onto polyvinylidene difluoride membrane (Amersham, GE Healthcare). The recombinant enzymes were probed using anti-V5-HRP conjugated antibody (Invitrogen), which was detected using an ECL Advance Western Blotting Detection Kit (Amersham) by a FluorChem SP imager (Alpha Innotech Corp., San Leandro, CA, U.S.A.). The band densities were semi-quantified with ImageJ software (Schneider *et al*., 2012).

### Lipid analysis

The yeast neutral lipid content was analyzed by the Nile red fluorescence assay as described previously (Xu *et al*., 2017). In brief, an aliquot (100 *µ*L) of yeast culture was incubated with 5 *µ*L of Nile red solution (0.1 mg/mL in methanol) into 96-well solid black plates (Corning Inc.). The fluorescence was measured before and after the addition of the Nile red solution with excitation and emission at 485 and 538 nm, respectively, using a Synergy H4 Hybrid reader (Biotek). The neutral lipid content in yeast was represented by the Nile red values which were calculated based on the change in fluorescence over OD_600_ (ΔF/OD_600_).

The TAG content and fatty acid composition of yeast cells and *N. benthamiana* leaf samples were analyzed using GC/MS. Yeast total lipids were extracted from approximately 30 mg of lyophilized yeast cells as described previously (Xu *et al*., 2019; Xu, Holic, *et al*., 2018). As for *N. benthamiana* leaf samples, 70 mg lyophilized leaf tissues were homogenized in chloroform: isopropanol (2:1, v/v) for total lipids extraction as described previously (Mietkiewska *et al*., 2014). For quantification, 100 *µ*g of triheptadecanoin (C17:0 TAG) were added to each sample as an internal standard. The extracted lipids were further separated on a TLC plate (0.25 mm Silica gel, DC-Fertigplatten) as described above and the lipid bands were visualized by spraying with primulin solution [0.05% primulin (w/v) in acetone/water (80:20, v/v)]. The corresponding TAG bands were then scraped and trans-methylated by incubating with 1 mL of 3 N methanolic HCl at 80 °C for 1 h. The resulting fatty acid methyl esters were analyzed using GC/MS (Agilent Technologies, Wilmington, DE) equipped with a capillary DB-23 column (30 m × 0.25 mm × 0.25 μm) as described previously (Xu *et al*., 2019; Xu, Holic, *et al*., 2018).

### Statistical analysis

Data are shown as means ± standard derivation (S.D.) for the number of independent experiments indicated. Significant differences between two groups were assessed using a Student’s t-test with the SPSS statistical package (SPSS 16.0, Chicago, IL, USA). The equality of variance was tested by Levene’s test. The unpaired Student’s t-test assuming equal variances and the unpaired Student’s t-test with Welch corrections assuming unequal variances were performed when the variances were equal and unequal, respectively.

## ACCESSION NUMBERS

Sequence data from this article can be accessed in the Phytozome/GenBank/Arabidopsis Genome Initiative databases under the following accession numbers: AtACBP6, NM_102916; AtGPAT9, AT5G60620; AtWRI1, AY254038; CzDGAT1, Cz09g08290.

## ACKNOWLEDGEMENTS

We acknowledge the support provided by the University of Alberta Start-up Research Grant, Natural Sciences and Engineering Research Council of Canada Discovery Grant (RGPIN-2016-05926) and the Canada Research Chairs Program. We are grateful to Dr. Mee-Len Chye (University of Hong Kong) for providing the plasmid pAT332 containing the coding sequence of *AtACBP6* and Dr. Igor Stagljar (University of Toronto) for providing the membrane yeast two-hybrid system. We also thank Dr. Stacy Singer (Agriculture and Agri-Food Canada) for sharing her experience on tobacco leaf infiltration, Dr. Michael Gänzle, Dr. Yuan Fang and Mr. Kosala Waduthanthri (University of Alberta) for their assistance in fluorescence microscopy and Dr. Shanjida Khan (University of Alberta) for providing the *N. benthamiana* seeds.

## AUTHOR CONTRIBUTIONS

GC supervised the experiment; YX, GC, and KMPC designed the experiment; YX performed the experiments and prepared the initial draft of the manuscript. YX, KMPC and LF analyzed the data. KMPC and KJ contributed valuable discussion during this study. All authors were involved in further editing of the manuscript.

## SUPPORTING INFORMATION

**Method S1.** Membrane yeast two-hybrid assay.

**Table S1.** DGAT1 proteins used for multiple sequence alignment.

**Table S2.** Apparent kinetic parameters of CzDGAT1 variants using a combined model accounting for sigmoidicity and substrate inhibition (Xu et al., 2017).

**Table S3.** Primers used in the current study.

**Figure S1.** Alignment of DGAT1 from different species.

**Figure S2.** Prediction of intrinsic disorder profile (blue) of the N-terminal region of DGAT1 from representative algae, plant and animals and its likelihood to participate in protein-protein interaction (red).

**Figure S3.** DGAT activity of CzDGAT1 variant enzymes at high oleoyl-CoA concentrations.

**Figure S4.** Probing possible self-interaction of CzDGAT1 variants using membrane yeast two-hybrid assay.

**Figure S5.** Enzyme activity of CzDGAT1 variants in the presence of Coenzyme A (CoA).

**Figure S6.** Illustration of the N-terminal fusion of acyl-CoA binding protein (ACBP) to CzDGAT1.

