## Supplementary files for "Kinetic improvement of an algal diacylglycerol acyltransferase 1 via fusion with an acyl-CoA binding protein"

**SUPPORTING INFORMATION**

**Method S1. Membrane yeast two-hybrid assay.**

**Table S1. DGAT1 proteins used for multiple sequence alignment.** The topology organization of DGAT1 was predicted using TMHMM (Krogh *et al.*, 2001). TMD, transmembrane domain.

**Table S2. Apparent kinetic parameters of CzDGAT1 variants using a combined model accounting for sigmoidicity and substrate inhibition** (Xu *et al.*, 2017)**.** DGAT activity was examined at increasing oleoyl-CoA concentration from 0.1 to 25 μM. Data were fitted to a nonlinear regression using a combined model accounting for sigmoidicity and substrate inhibition with the GraphPad Prism software. Data shown are means ± S.D. (n=3).

**Table S3. Primers used in the current study.** A restriction site or linker is shown in **bold and underlined** and a Kozak translation initiation sequence for yeast expression is shown in *italic*.

**Figure S1**. **Alignment of DGAT1 from different species.** Alignment was virtualized by Geneious v5.3 (Drummond *et al.*, 2010).

**Figure S2**. **Prediction of intrinsic disorder profile (blue) of the N-terminal region of DGAT1 from representative algae, plant and animals and its likelihood to participate in protein-protein interaction (red).** The analyses were performed by DISOPRED (Ward *et al.*, 2004). *At*, *Arabidopsis thaliana*; *Kn*, *Klebsormidium nitens*; *Cz*, *Chromochloris zofingiensis*; *Pt*, *Phaeodactylum tricornutum*; *Tp*, *Thalassiosira pseudonana*.

**Figure S3. DGAT activity of CzDGAT1 variant enzymes at high oleoyl-CoA concentrations.** A-D, DGAT activities of the full-length CzDGAT1 ($DGAT1_{1-550}$), N-terminal truncated CzDGAT1 (DGAT1$_{81-550}$) and their corresponding acyl-CoA binding protein (ACBP) fused proteins (ACBP-DGAT1$_{1-550}$ and ACBP-DGAT1$_{81-550}$) at oleoyl-CoA concentration from 0.1-25 μM. Data were fitted to the allosteric sigmoidal equation or a previously proposed kinetic model that accounts for sigmoidicity and substrate inhibition (Xu *et al.*, 2017) using GraphPad Prism. The combined kinetic model is the preferred model for CzDGAT1$_{1-550}$, ACBP-fused CzDGAT1$_{1-550}$, and ACBP-fused CzDGAT1$_{81-550}$, but not CzDGAT1$_{81-550}$. Data represent means ± S.D. (n = 3).

**Figure S4. Probing possible self-interaction of CzDGAT1 variants using membrane yeast two-hybrid assay.** DNA sequences encoding CzDGAT1$_{1-550}$, CzDGAT1$_{81-550}$ and CzDGAT1$_{107-550}$ were ligated to the Lex A- C-terminal fragment of ubiquitin ($C_{ub}$) and the N-terminal fragment of ubiquitin containing an Ile/Gly point mutation ($N_{ub}G$), yielding $C_{ub}$-bait and $N_{ub}G$-prey, respectively. Serial dilutions of yeast cells producing each bait/prey combination were spotted on synthetic drop-out (SD) agar plates lacking Ade, His, Leu and Trp (SD-A-H-L-T).

**Figure S5. Enzyme activity of CzDGAT1 variants in the presence of Coenzyme A (CoA).** The DGAT activities of CzDGAT1$_{1-550}$, and acyl-CoA binding protein (ACBP) fused enzyme (ACBP-DGAT1$_{1-550}$) were assayed at 5 μM oleoyl-CoA in the absence or presence of 50 μM CoA. Data represent means ± S.D. (n = 3).

**Figure S6. Illustration of the N-terminal fusion of acyl-CoA binding protein (ACBP) to CzDGAT1.** The three-dimensional structure of *Arabidopsis thaliana* ACBP6 was generated with the SWISS-MODEL software. *A. thaliana* ACBP6 consists of four α helixes for acyl-CoA binding and is proposed to facilitate the feeding of acyl-CoA to the catalytic pocket of CzDGAT1 via capturing cytosolic acyl-CoAs or acyl-CoAs partitioned into the membrane lipid bilayer and subsequently channeling them to DGAT by proximity.

**Method S1. Membrane yeast two-hybrid assay.**

Possible self-interaction of $CzDGAT1_{1-550}$ and its N-terminal truncated mutants ($CzDGAT1_{81-550}$ and $CzDGAT1_{107-550}$) were tested using the membrane yeast two-hybrid system (kindly provided by Dr. Igor Stagljar, University of Toronto) with the method described by Snider et al. (Snider *et al.*, 2010). Briefly, cDNAs encoding $CzDGAT1_{1-550}$, $CzDGAT1_{81-550}$ and $CzDGAT1_{107-550}$ were amplified by PCR and cloned into the pBT3N bait vector or pPR3N prey vector, respectively. The pBT3N:bait was then co-transformed with the pPR3N:prey, Ost-$N_{ub}$I 'positive' control prey or Ost-$N_{ub}$G 'negative' control prey into the yeast strain NMY51 [*MATa, his3Δ200, trp1-901, leu2-3,112, ade2, LYS2::(lexAop)4-HIS3,ura3::(lexAop)8-lacZ,ade2::(lexAop)8-ADE2, GAL4*]. The possible interaction was then assayed on synthetic drop-out (SD) agar plates lacking Ade, His, Leu and Trp (SD-A-H-L-T) by 1: 10 serial dilution of cell cultures starting from an OD600 value of 0.4.

**Table S1. DGAT1 proteins used for multiple sequence alignment.** The topology organization of DGAT1 was predicted using TMHMM (Krogh *et al.*, 2001). TMD, transmembrane domain.

| Name | Organism | Phytozome/Genbank accession number/JGI protein ID | Length of N-terminus (amino acid residues) | # of TMD |
|---|---|---|---|---|
| ApDGAT1 | *Auxenochlorella protothecoides* | XP_011402032 | 326 | 7 |
| AtDGAT1 | *Arabidopsis thaliana* | NM_127503 | 132 | 9 |
| BnDGAT1 | *Brassica napus* | JN224473 | 113 | 8 |
| BtDGAT1 | *Bos taurus* | AAL49962 | 83 | 9 |
| CaeDGAT1 | *Caenorhabditis elegans* | NM_001269372 | 70 | 9 |
| CheDGAT1 | *Chlorella ellipsoidea* | KT779429 | 260 | 9 |
| CreDGAT1 | *Chlamydomonas reinhardtii* | Cre01.g045903 | 158 | 7 |
| CsDGAT1 | *Camelina sativa* | XM_010417066 | 132 | 9 |
| CsuDGAT1 | *Coccomyxa subellipsoidea C-169* | 54084 | 1 | 9 |
| CvDGAT1 | *Chlorella vulgaris* | ALP13863.1 | 20 | 9 |
| CzDGAT1A | *Chromochloris zofingiensis* | MH523419 | 289 | 9 |
| CzDGAT1B | *Chromochloris zofingiensis* | Cz09g08290 | 107 | 9 |
| DmDGAT1 | *Drosophila melanogaster* | AF468649 | 132 | 8 |
| DrDGAT1 | *Danio rerio* | NM_199730 | 86 | 9 |
| EaDGAT1 | *Euonymus alatus* | AY751297 | 111 | 9 |
| GmDGAT1 | *Glycine max* | AY496439 | 102 | 9 |
| HaDGAT1 | *Helianthus annuus* | HM015632 | 110 | 9 |
| HsDGAT1 | *Homo sapiens* | NM_012079 | 86 | 9 |
| JcDGAT1 | *Jatropha curcas* | DQ278448 | 123 | 9 |
| KnDGAT1 | *Klebsormidium nitens* | GAQ91878 | 260 | 9 |
| LuDGAT1 | *Linum usitatissimum* | KC485337 | 111 | 9 |
| MdDGAT1 | *Monodelphis domestica* | XM_007488766 | 227 | 8 |
| MmDGAT1 | *Mus musculus* | AF078752 | 95 | 9 |
| MtDGAT1 | *Medicago truncatula* | XM_003595183 | 142 | 9 |
| NoDGAT1 | *Nannochloropsis oceanica* | KY073295 | 29 | 9 |
| NtDGAT1 | *Nicotiana tabacum* | AF129003 | 140 | 9 |
| NvDGAT1 | *Nematostella vectensis* | XM_001639301 | 54 | 9 |
| OeDGAT1 | *Olea europae* | AY445635 | 136 | 9 |
| OsDGAT1 | *Oryza sativa* | NM_001061404 | 142 | 8 |
| PbDGAT1 | *Paracoccidioides brasiliensis* | EEH17170.1 | 79 | 10 |
| PfDGAT1 | *Perilla frutescens* | AF298815 | 138 | 9 |
| PotDGAT1 | *Populus trichocarpa* | XM_006371934 | 96 | 9 |
| PpDGAT1 | *Physcomitrella patens* | XM_001770877 | 51 | 9 |
| PtDGAT1 | *Phaeodactylum tricornutum* | HQ589265 | 149 | 8 |

| | | | | |
|---|---|---|---|---|
| RcDGAT1 | *Ricinus communis* | NM_001323734 | 128 | 9 |
| RnDGAT1 | *Rattus norvegicus* | AB062759 | 97 | 9 |
| SiDGAT1 | *Sesamum indicum* | JF499689 | 147 | 9 |
| SsDGAT1 | *Sus scrofa* | NM_214051 | 83 | 9 |
| TaDGAT1 | *Trichoplax adhaerens* | XM_002111989 | 11 | 9 |
| TgDGAT1 | *Toxoplasma gondii* | AY327327 | 117 | 9 |
| TmDGAT1 | *Tropaeolum majus* | AY084052 | 124 | 9 |
| TpDGAT1 | *Thalassiosira pseudonana* | XM_002287179 | 38 | 9 |
| VfDGAT1 | *Vernicia fordii* | DQ356680 | 129 | 9 |
| VgDGAT1 | *Vernonia galamensis* | EF653276 | 127 | 9 |
| VvDGAT1 | *Vitis vinifera* | CAN80418 | 103 | 9 |
| YlDGAT1 | *Yarrowia lipolytica* | XM_502557 | 98 | 8 |
| ZmDGAT1 | *Zea mays* | EU039830 | 97 | 8 |

**Table S2. Apparent kinetic parameters of CzDGAT1 variants using a combined model accounting for sigmoidicity and substrate inhibition** (Xu *et al.*, 2017)**.** DGAT activity was examined at increasing oleoyl-CoA concentration from 0.1 to 25 μM. Data were fitted to a nonlinear regression using a combined model accounting for sigmoidicity and substrate inhibition with the GraphPad Prism software. Data shown are means ± S.D. (n=3).

| Enzyme | $DGAT1_{1\text{-}550}$ | $DGAT1_{81\text{-}550}$ | ACBP-$DGAT1_{1\text{-}550}$ | ACBP-$DGAT1_{81\text{-}550}$ |
|---|---|---|---|---|
| Preferred model | Combined model | Allosteric sigmoidal | Combined model | Combined model |
| Apparent $V_{max}$ (pmol TAG/min/mg protein) | 314.20±16.11 | 16.82±0.17 | 502.50±23.75 | 79.38±27.04 |
| Hill coefficient | 1.31±0.08 | 1.76±0.06 | 1.72±0.12 | 1.12±0.14 |
| Apparent $S_{0.5}$ (μM) | 1.76±0.15 | 1.78±0.04 | 1.10±0.08 | 4.76±2.52 |
| Apparent $K_i$ (μM) | 58.74±11.85 | - | 38.44±7.03 | 9.03±4.41 |
| Goodness of Fit/R2 | 0.992 | 0.995 | 0.980 | 0.961 |

**Table S3. Primers used in the current study.** A restriction site or linker is shown in **bold and underlined** and a Kozak translation initiation sequence for yeast expression is shown in *italic*.

| Primer name | Oligonucleotide sequence |
|---|---|
| Primers used for cloning of cDNAs for yeast expression | |
| CzDGAT1$_{1-550}$-F (to pYES2.1) | gcaga**<u>GCGGCCGC</u>***GAA*ATGGAGGGTGCACGAATC |
| CzDGAT1-R (to pYES2.1) | tat**<u>GTCGAC</u>**GTGTGACATAAGAGCAGCATTCC |
| CzDGAT1$_{81-550}$-F (to pYES2.1) | gcaga**<u>GCGGCCGC</u>***GAA*ATGATTGGCAGCAGTCCTCT |
| CzDGAT1$_{107-550}$-F (to pYES2.1) | gcaga**<u>GCGGCCGC</u>***GAA*ATGGATGGACTGGTCAACTTGGC |
| AtACBP6-F (to pYES2.1) | gcaga**<u>GCGGCCGC</u>***GAA*ATGGGTTTGAAGGAGGAATTTG |
| AtACBP6-R (to pYES2.1) | tat**<u>GTCGAC</u>**GGTTGAAGCCTTGGAAGCA |
| AtACBP6-Linker-R (to pYES2.1) | **<u>GAATTCCGT</u>**GGTTGAAGCCTTGGAAGCA |
| Linker-CzDGAT1$_{1-550}$-F (to pYES2.1) | **<u>ACGGAATTC</u>**ATGGAGGGTGCACGAATC |
| Linker-CzDGAT1$_{81-550}$-F (to pYES2.1) | **<u>ACGGAATTC</u>**ATGATTGGCAGCAGTCCTCT |
| Linker-CzDGAT1$_{107-550}$-F (to pYES2.1) | **<u>ACGGAATTC</u>**ATGGATGGACTGGTCAACTTGGC |
| AtACBP6-F (to pESC-leu2d empty) | tata**<u>GGATCC</u>***GAA*ATGGGTTTGAAGGAGGAATTTG |
| AtACBP6-R (to pESC-leu2d empty) | tata**<u>CTCGAG</u>**TCAGGTTGAAGCCTTGGAAGCA |
| Primers used for cloning of cDNAs for membrane yeast two-hybrid assay (MYTH) | |
| CzDGAT1$_{1-550}$-F (to MYTH vector) | ac**<u>GGCCATTACGGCC</u>**ATGGAGGGTGCACGAATC |
| CzDGAT1$_{81-550}$-F (to MYTH vector) | ac**<u>GGCCATTACGGCC</u>**ATGATTGGCAGCAGTCCTCT |
| CzDGAT1$_{107-550}$-F (to MYTH vector) | ac**<u>GGCCATTACGGCC</u>**ATGGATGGACTGGTCAACTTGGC |
| CzDGAT1-R (to MYTH vector) | tat**<u>GGCCGAGGCGGCC</u>**TCAGTGTGACATAAGAGCAGCATTCC |
| Primers used for cloning of cDNAs for plant expression | |
| AtWRI-F (to pGreen) | CTAG**<u>ACTAGT</u>**ATGAAGAAGCGCTTAACCAC |
| AtWRI-R (to pGreen) | CGC**<u>TCTAGA</u>**TCAGACCAAATAGTTACAAGAAAC |
| AtACBP-F (to pGreen) | TATA**<u>ACCGGT</u>**ATGGGTTTGAAGGAGGAATTTG |
| AtACBP-R (to pGreen) | CGC**<u>TCTAGA</u>**TCAGGTTGAAGCCTTGGAAGCA |
| CzDGAT1 $_{1-550}$-F (to pGreen) | TATA**<u>ACCGGTACTAGT</u>**ATGGAGGGTGCACGAATC |
| CzDGAT1 $_{81-550}$-F (to pGreen) | TATA**<u>ACCGGT</u>**ATGATTGGCAGCAGTCCTCT |
| CzDGAT1 $_{107-550}$-F (to pGreen) | TATA**<u>ACCGGT</u>**ATGGATGGACTGGTCAACTTGGC |
| CzDGAT1-R (to pGreen) | CGC**<u>TCTAGA</u>**TCAGTGTGACATAAGAGCAGCATTCC |
| Venus-F (to pGPTVII) | cgc**<u>TCTAGA</u>**ATGGTGAGCAAGGGCGA |
| Venus-R (to pGPTVII) | TATA**<u>ACTAGTACCGGT</u>**CTTGTACAGCTCGTCCAT |
| AtACBP-R (to pGPTVII) | tata**<u>CTCGAG</u>**TCAGGTTGAAGCCTTGGAAGCA |
| CzDGAT1-R (to pGPTVII) | tata**<u>CTCGAG</u>**TCAGTGTGACATAAGAGCAGCATTCC |
| Linker-SCFP3A-F (to pGPTVII) | **<u>ACGGAATTC</u>**ATGGTGAGCAAGGGCGAGG |
| SCFP3A-R (to pGPTVII) | TATA**<u>CTCGAG</u>**TTACTTGTACAGCTCGTCCATGCC |
| AtGPAT9-F (to pGPTVII) | CGC**<u>TCTAGAGTCGAC</u>**ATGAGCAGTACGGCAGGGAG |
| AtGPAT9-linker-R (to pGPTVII) | **<u>GAATTCCGT</u>**CTTCTCTTCCAATCTAGCCAGGA |

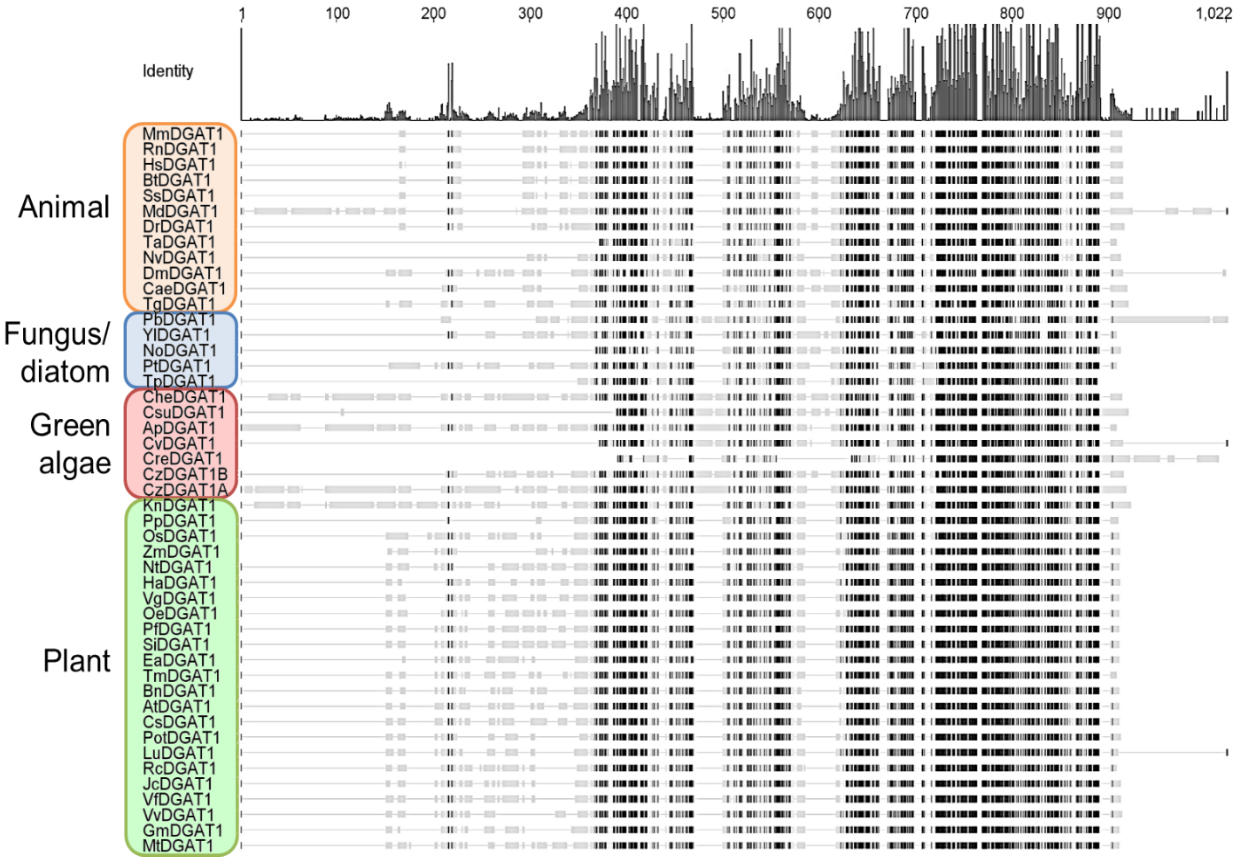


**Figure S1. Alignment of DGAT1 from different species.** Alignment was virtualized by Geneious v5.3 (Drummond *et al.*, 2010).

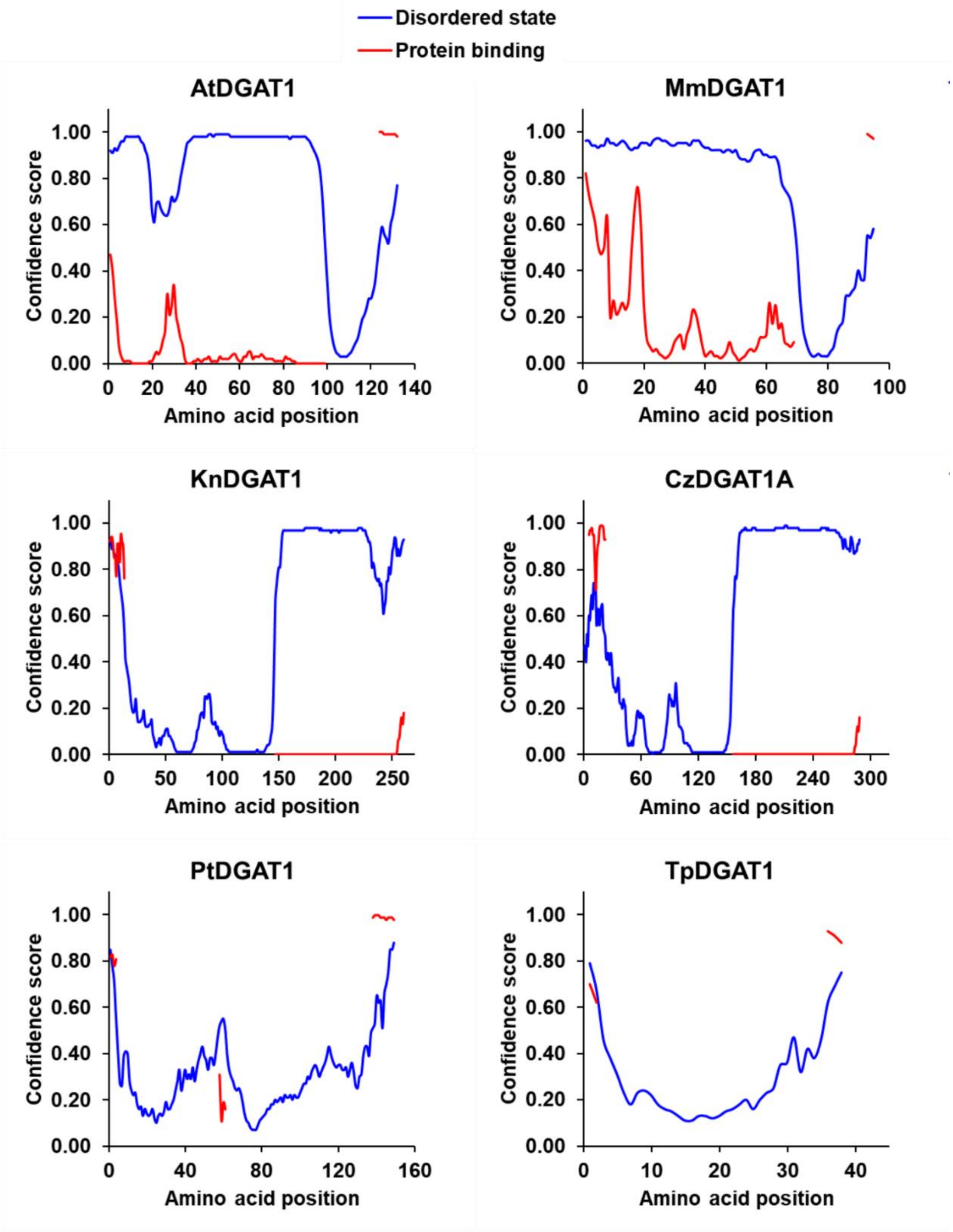


**Figure S2**. **Prediction of intrinsic disorder profile (blue) of the N-terminal region of DGAT1 from representative algae, plant and animals and its likelihood to participate in protein-protein interaction (red).** The analyses were performed by DISOPRED (Ward *et al.*, 2004). *At*, *Arabidopsis thaliana*; *Kn*, *Klebsormidium nitens*; *Cz*, *Chromochloris zofingiensis*; *Pt*, *Phaeodactylum tricornutum*; *Tp*, *Thalassiosira pseudonana*.

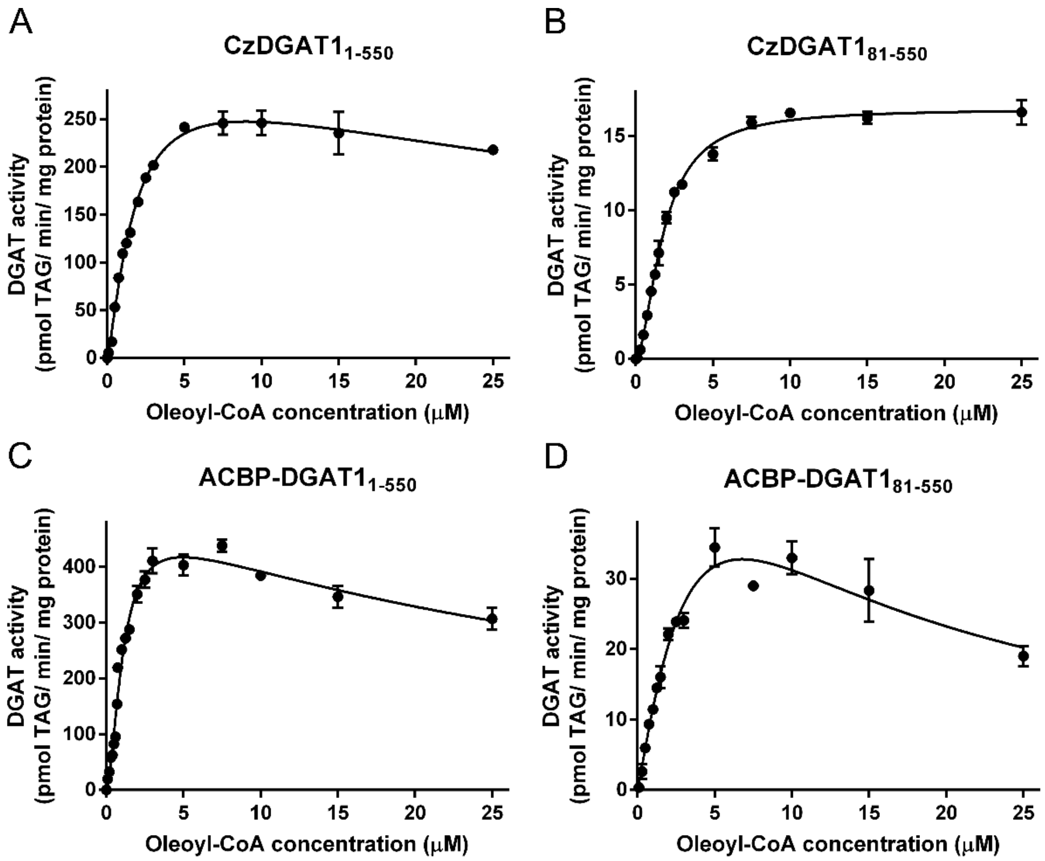


**Figure S3. DGAT activity of CzDGAT1 variant enzymes at high oleoyl-CoA concentrations.** A-D, DGAT activities of the full-length CzDGAT1 ($DGAT1_{1-550}$), N-terminal truncated CzDGAT1 ($DGAT1_{81-550}$) and their corresponding acyl-CoA binding protein (ACBP) fused proteins ($ACBP\text{-}DGAT1_{1-550}$ and $ACBP\text{-}DGAT1_{81-550}$) at oleoyl-CoA concentration from 0.1-25 µM. Data were fitted to the allosteric sigmoidal equation or a previously proposed kinetic model that accounts for sigmoidicity and substrate inhibition (Xu *et al.*, 2017) using GraphPad Prism. The combined kinetic model is the preferred model for $CzDGAT1_{1-550}$, ACBP-fused $CzDGAT1_{1-550}$, and ACBP-fused $CzDGAT1_{81-550}$, but not $CzDGAT1_{81-550}$. Data represent means ± S.D. (n = 3).

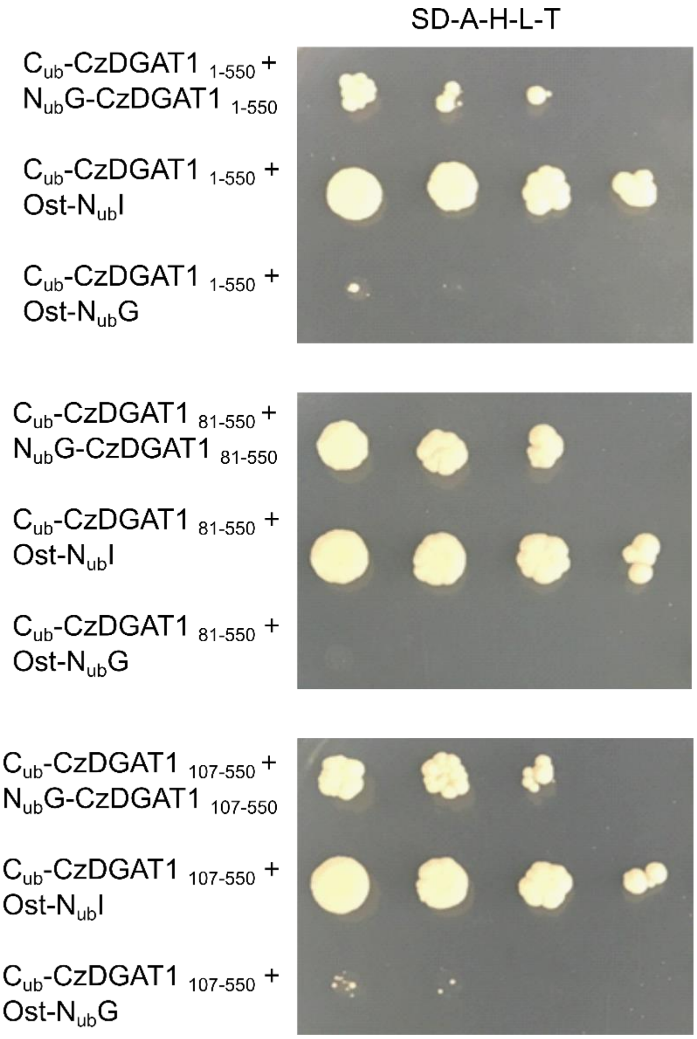


**Figure S4. Probing possible self-interaction of CzDGAT1 variants using membrane yeast two-hybrid assay.** DNA sequences encoding $CzDGAT1_{1\text{-}550}$, $CzDGAT1_{81\text{-}550}$ and $CzDGAT1_{107\text{-}550}$ were ligated to the Lex A- C-terminal fragment of ubiquitin ($C_{ub}$) and the N-terminal fragment of ubiquitin containing an Ile/Gly point mutation ($N_{ub}G$), yielding $C_{ub}$-bait and $N_{ub}G$-prey, respectively. Serial dilutions of yeast cells producing each bait/prey combination were spotted on synthetic drop-out (SD) agar plates lacking Ade, His, Leu and Trp (SD-A-H-L-T).

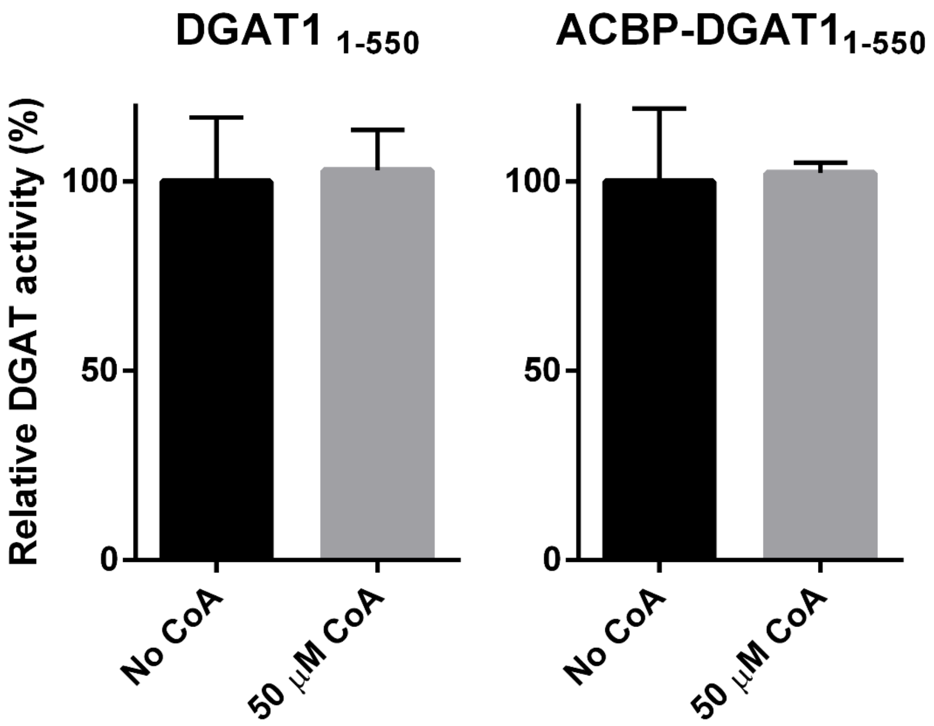


**Figure S5. Enzyme activity of CzDGAT1 variants in the presence of Coenzyme A (CoA).** The DGAT activities of CzDGAT1$_{1\text{-}550}$, and acyl-CoA binding protein (ACBP) fused enzyme (ACBP-DGAT1$_{1\text{-}550}$) were assayed at 5 μM oleoyl-CoA in the absence or presence of 50 μM CoA. Data represent means ± S.D. (n = 3).

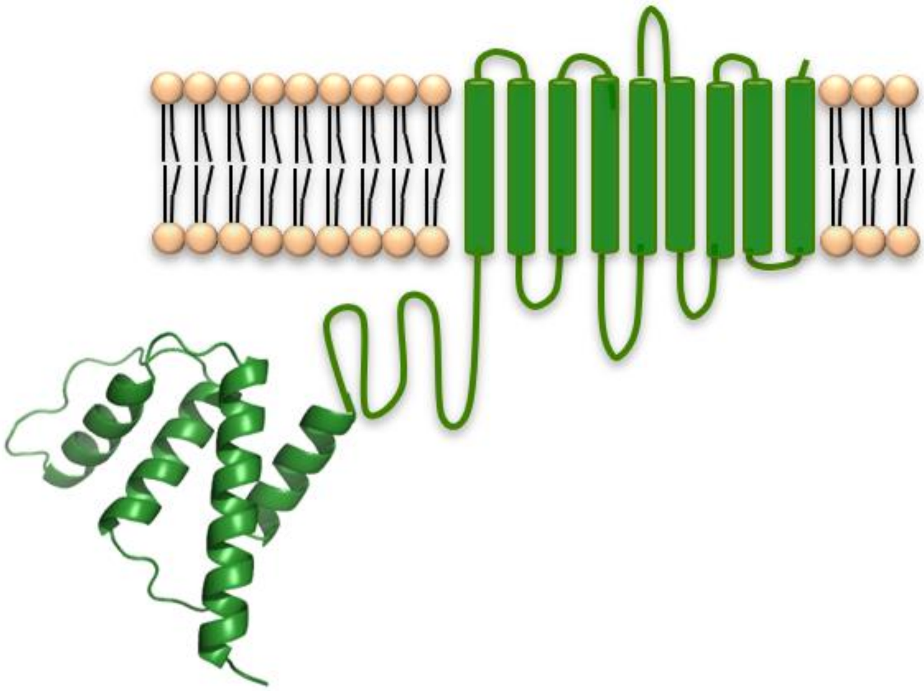

**Figure S6. Illustration of the N-terminal fusion of acyl-CoA binding protein (ACBP) to CzDGAT1.** The three-dimensional structure of *Arabidopsis thaliana* ACBP6 was generated with the SWISS-MODEL software. *A. thaliana* ACBP6 consists of four α helixes for acyl-CoA binding and is proposed to facilitate the feeding of acyl-CoA to the catalytic pocket of CzDGAT1 via capturing cytosolic acyl-CoAs or acyl-CoAs partitioned into the membrane lipid bilayer and subsequently channeling them to DGAT by proximity.
